# AI-assisted imaging screening reveals mechano-molecular tissue organizers and network of signaling hubs

**DOI:** 10.1101/2024.11.14.623670

**Authors:** Cristina Bertocchi, Juan José Alegría, Sebastián Vásquez-Sepúlveda, Rosario Ibanez-Prat, Aishwarya Srinivasan, Ignacio Arrano-Valenzuela, Barbara Castro-Pereira, Catalina Soto-Montandon, Alejandra Trujillo-Espergel, Gareth I. Owen, Pakorn Kanchanawong, Mauricio Cerda, Giovanni Motta, Ronen Zaidel-Bar, Andrea Ravasio

## Abstract

Cadherin-mediated adhesions are crucial mechanical and signaling hubs that connect cells within a tissue and probe the mechanics of the surrounding environment. They constitute a physical link between the actin cytoskeleton of neighboring cells, providing the mechanical coordination needed for morphogenetic processes, tissue homeostasis, collective migration, and regeneration. Disruptions in adhesion mechanisms are closely linked to the breakdown of epithelial structure and the emergence of disease-related traits characteristic of cancer progression. The cadhesome network comprises over 170 structural and regulatory proteins involved in cadherin-mediated adhesion. While this network is essential for coordinating tissue responses to mechanical stress, its complexity has historically limited our understanding of how individual components contribute to force transmission and tissue homeostasis. Recent technological advances offer tools to investigate large molecular networks in cellular function and pathology (functional omics). Leveraging these advances, we developed an experimental and analytical platform combining high-throughput gene silencing, imaging, and artificial intelligence (AI) to systematically profile each role of each protein in tissue formation, mechanical stability, and response to induced tension. Using EpH4 cells as an epithelial tissue model, we performed systematic silencing in triplicate, capturing a range of tissue phenotypes under baseline and tension-inducing conditions. Machine learning methods were used to analyze complex imaging data, quantify tissue ruptures, characterize junctional organization, and measure tension states of the tissue. By incorporating machine learning algorithms, we automated image feature extraction, clustering, and classification, enabling an unprecedented quantitative evaluation of tissue mechanics at scale. Our machine learning models allowed us to identify significant patterns, including protein-specific responses to tension and their roles in tissue-level mechanical integrity. Finally, we constructed a protein interaction network detailing the roles of each protein, their physical interactions, and known links to cancer. The network analysis revealed three prominent mechanotransductive and signaling subnetworks centered around E-cadherin, EGFR, and RAC1. Our study provides a foundational framework for investigating mechanosensing proteins and it offers a scalable blueprint for discovering potential therapeutic targets in diseases like cancer, where tissue mechanics play a crucial role.

**Teaser:** AI-aided screening identifies key regulators of epithelial tissue mechanics, uncovering potential therapeutic targets in cancer.

## Introduction

The evolution of multicellularity coincides with the emergence of conserved molecular complexes that mediate dynamic connections between cells, enabling tissues to establish physiological function and to withstand and respond to mechanical forces ^1^. The ability of these protein complexes to sustain tissue homeostasis through cell-to-cell force transmission relies on the precise, hierarchical organization of their molecular components into a network that collectively integrates mechanical and biochemical signals ^2–4^. This functional integration is essential for regulating intercellular adhesions, mechanical adaptation, and mechanotransduction ^5^. Among these complexes, cadherin-mediated adhesion serves as crucial hub, linking the actin cytoskeleton of neighboring cells to maintain mechanical coordination required for processes such as morphogenesis, tissue homeostasis, collective migration, and tissue regeneration ^6^. Consistently, defects in cell-cell adhesion are strongly associated with the loss of organized epithelial structure and the development of diseases-promoting features, such as uncontrolled cell growth, survival, and increased tendencies for cell migration and invasion – hallmarks commonly associated with cancerous transformation and are pivotal in driving tumor progression ^7^. The expanded network of cadherin-associated proteins that mediate and regulate cell-cell adhesion is known as the "cadhesome" ^8^. This extensive network includes over 170 structural and regulatory proteins that connect transmembrane cadherins to the actin cytoskeleton, coordinating dynamic responses within tissues. The richness and complexity of the cadhesome have posed challenges to fully understanding how individual proteins contribute to mechanical regulation and force transmission. Consequently, while significant research has been conducted across various scales to reveal the molecular mechanisms of force transmission and its role in maintaining tissue homeostasis, understanding the network from a system level complexity remains elusive^9^, thus limiting our knowledge of how the cadhesome drives tissue formation, mechanical stability and response to mechanical tension.

Recent technological advances in experimental methods to generically manipulate cells, high-throughput imaging techniques and advanced statistical and computational analysis paved the way to explore functional complexity of biological systems through functional omics^10^. As an emerging frontier in biological investigation, functional omics systematically examines the roles and interactions of molecules to understand their effects on cellular functions and phenotypes, offering a framework for linking protein networks to cellular behaviors and biomedically-relevant pathological processes. To harness this potential, we developed an experimental and analytical strategy that combines gene silencing, high-throughput imaging, and unbiased image analysis to screen and categorize each component of the cadhesome based on its role in tissue formation, mechanical homeostasis, and response to pharmacologically induced tension. Additionally, we utilized online databases and tools to construct a network detailing the mechanical roles of each protein, their physical interactions, and the reported correlation between gene expression and breast cancer patient survival. Our findings offer a foundational blueprint for identifying novel mechanosensing proteins that regulate tissue tensional homeostasis. The approach developed here provides a robust framework for systematically exploring the molecular mechanisms underlying complex biological functions and could serves as a guideline for designing similar experimental strategies in functional omics.

## Results

### Design of high-throughput siRNA screening of proteins of the cadhesome network

EpH4 murine cells were chosen as the experimental model of an epithelial tissue due to their amenability to genetic manipulation. As a first step, we checked the expression levels of cadhesome genes in EpH4 cell by RNA seq technology (Suppl. Table 1). This analysis revealed that six genes were not expressed in EpH4 (AJAP1, CTNNA2, CTNNA3, LIN7A, LPP and PPP2CA) and thus they were excluded from further analysis. To probe the role of canonical cadhesome proteins in regulating mechanical functions of epithelial tissues (i.e. formation and maintenance of tissue integrity and mechanical homeostasis under low and high tension), we developed a high-throughput screen based on silencing gene expression (Figure 1A and Suppl. Figure 1). EpH4 cells were transfected in suspension with SMART pools of siRNAs to individually silence each one of the 167 bona fide components of the cadhesome (samples called GeneX-siRNA in Figure 1B where X stands for a particular cadhesome protein), while non-target siRNA was used to generate reference wildtype tissues under the same transfection conditions (samples Neg-siRNA in Figure 1B). Since each protein silencing was performed in triplicate and on different days, thirty-four 96-well plates were used to accommodate all of the samples. Validation of siRNA silencing was performed by Western Blot on cells transfected with siRNA SMARTpool targeting CDH1 (Suppl. Figure 2A). Transfected cells were seeded in 96-well plates and allowed to grow for 24 hours to form a confluent monolayer. Aside from samples with silenced cadhesome proteins (GeneX-siRNA), each 96-well plate also contained Neg-siRNA samples and controls for transfection efficiency (siGLO and siTOX – Suppl. Figure 2B). Imaging and analysis of the 96-well plates was conducted only if visual inspection of siGLO and siTOX samples confirmed successful transfection. siRNA transfected samples were seeded in duplicates to conduct paired observation of tissue at normal tensional state (samples labeled as Vehicle in Figure 1B) and tissue with elevated tension (samples labeled as Nocodazole in Figure 1B), which was obtained by chemically stimulating cells by Nocodazole (5µM for 2 hours prior cell fixation). Use of nocodazole to pharmacologically stimulate tension in cell monolayers has been previously demonstrated by various groups, including ours ^3,11–14^. Here, we further validated the use of nocodazole as a mean to induce tension in EpH4 tissues by observing a noticeable increase in actin tensional elements, such as stress fibers, in nocodazole treated cells as compared to non-treated conditions (Suppl. Figure 3). Importantly, nocodazole treatment seems to elicit a larger cellular response when compared to mechanical stimulation performed on the same cell line (60% axial stretch, Suppl. Figure 4). 24 hours after seeding, samples were treated with vehicle or nocodazole for 2 hours, and thereafter fixed and stained for cell-cell adhesion using antibody against E-Cadherin and nuclei using DAPI. Additionally, randomly selected samples were stained for F-Actin (phalloidin) to qualitatively evaluate tissue tensional state by examining characteristic tension-associated actin structures (e.g., stress fibers). An automated image acquisition using Metamorph-based high-content screening (HCS) module was performed to acquire a matrix of adjacent z-stacks images that were z average projected and stitched to obtain a large field of view (3x3 images, corresponding to a total of 1489 by 1489 pixels).

**Figure 1.**
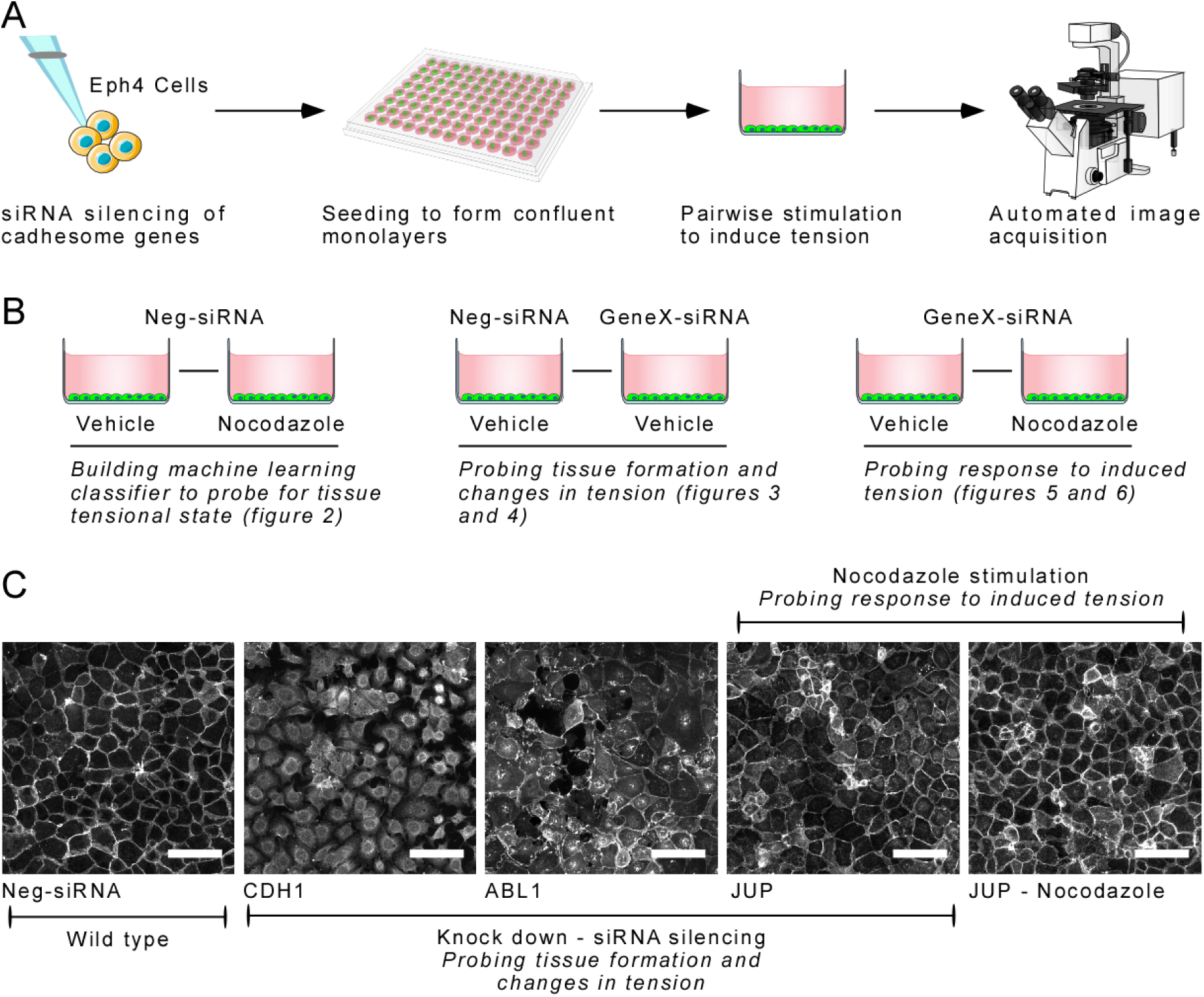
High-throughput screening to discover proteins involved in tissue mechanical regulation. A – Schematic representation of the high-throughput assay (see also suppl. Figure 1A). Eph4 cells were transfected in suspension by lipofectamine with mouse SMART pools siRNAs targeting bona fide components of the cadhesome (GeneX-siRNA) or a scrambled sequence (Neg-SiRNA) as control for wild type tissue. Cells were then seeded in glass-bottom 96-well plates and allowed to grow for 24 hours to form a monolayer. Treatment with nocodazole (5µM for 2 hours) was used as pharmacological stimulation to induce tension within the tissue (Suppl figure 2), whereas vehicle (DMSO) was used as control for non-stimulated tissue. Following fixation and permeabilization, cells were stained for E-Cadherin to visualize the cell-cell junctions. The resulting cell monolayers have been imaged using a CSU spinning disk (Yokogawa®) confocal head mounted on an inverted microscope (Eclipse Ti, Nikon, Japan). Images were acquired using a Plain Apo Vc 60X oil objective, N.A. 1.40, with 561nm excitation wavelengths for Alexa 568 (to detect the secondary antibody versus E-Cadherin antibody). High-content image acquisition of fixed cell was performed using a Metamorph based high-content screening (HCS) module. Nine adjacent z-stacks (step 0.5um, total height 3.5um) have been averaged to a projection and stitched in a 3x3 montage (Suppl. Figure 1A). B – Schematic of the resulting samples pairs. Each 96-well contained 2 wells transfected with scrambled siRNA (Neg-siRNA) to access the appearance of the wildtype monolayer (left pair). Within this pair, one was stimulated with Nocodazole to induce tension while the other one served as the vehicle-only control (Vehicle), allowing assessment of the effect of induced tension on the wild-type epithelial tissue. This sample pair was used to generate the classifier via machine learning (Figure 2). Cells in the remaining wells were transfected with siRNA to silence the cadhesomés bona fides proteins (GeneX-siRNA). For each protein, two wells were seeded: One well was treated with Nocodazole to induce tension while the other remained untreated (Vehicle). Untreated tissues silenced for a specific cadhesome protein (GeneX-siRNA) can be compared with untreated wild type tissues (Neg-siRNA) (central pair), allowing to evaluate the effect of specific protein silencing on the tensional state of the tissue (Figures 3, 4 and suppl. Figure 7). Finaly, tissues silenced for specific proteins (GeneX-siRNA) nocodazole-treated and non-treated (vehicle) sample (right pair) can be compared with those where tension was pharmacologically induced (nocodazole-treated) and non-treated (vehicle), allowing to evaluate the effect of specific protein silencing on tissue stability in response to tension induction (Figures 5, 6, and suppl. Figure 8). C – Exemplary images illustrating various cellular responses. From left to right: Cells transfected with scrambled RNA sequence (Neg-siRNA) displayed the characteristic appearance of unperturbed epithelial tissue (wild type). Silencing of E-Cadherin (CDH1) resulted in a complete rupture of the tissue, while silencing of Abl1 led to partial rupture. Tissue ruptures could also occur in some GeneX-siRNA samples only after nocodazole stimulation. Besides these extreme phenotypes leading to varying degrees of tissue rupture, most proteins silencing and/or nocodazole stimulation resulted in more subtle changes. An example of such cases is here provided (JUP). Untreated and nocodazole-treated tissues display only subtle differences between each other and when visually compared to the wild type. Thus, we resorted to the use of a machine learning classifier trained on untreated (vehicle) and nocodazole-treated Neg-siRNA samples to quantitatively compare such samples. Samples are referred to untreated condition (vehicle), if not otherwise indicated. Scale bar = 50μm.

**Figure 2.**
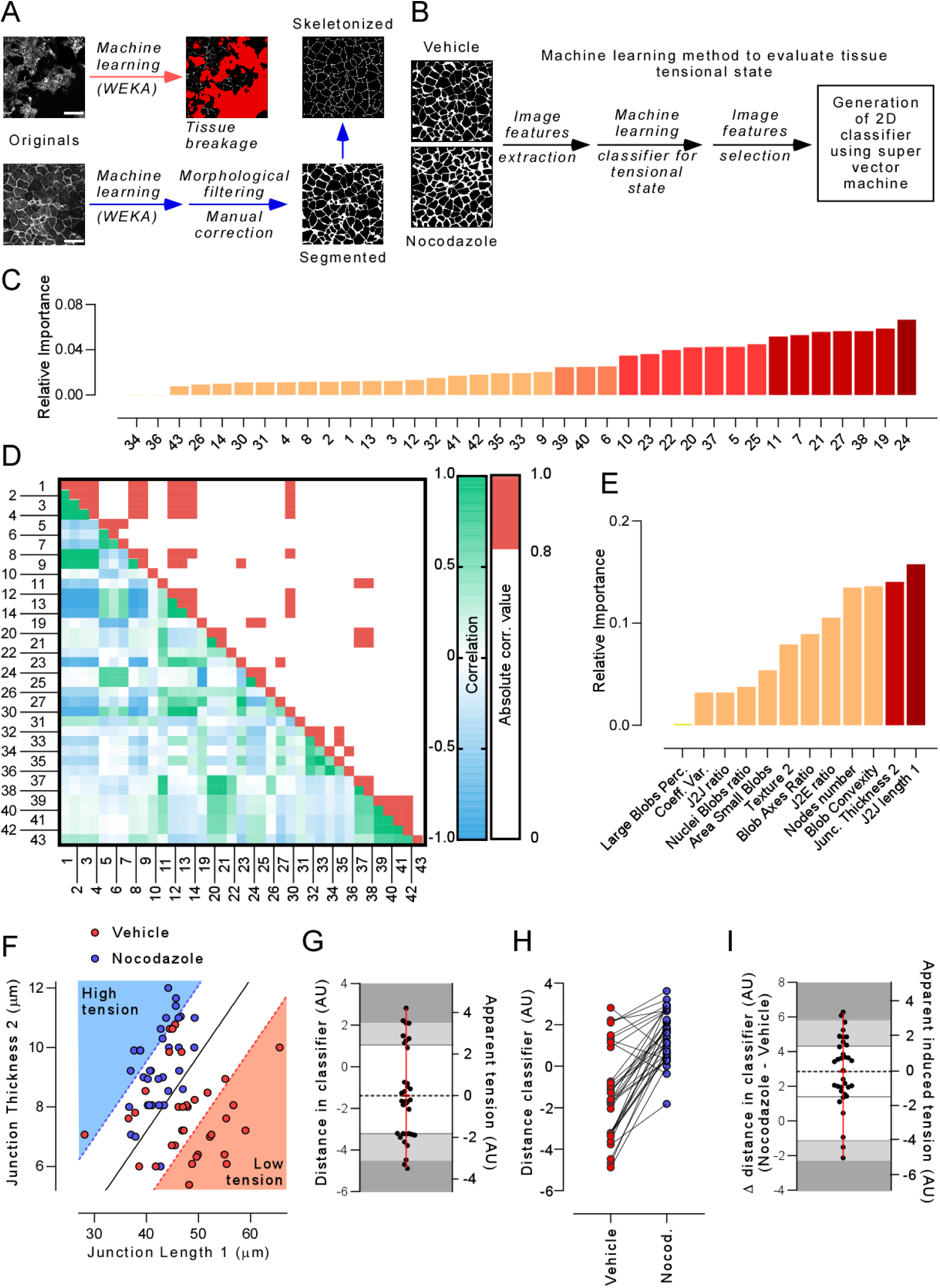
Classification Pipeline for Quantifying Tissue Tensional State. A – Description of the pipeline used for image segmentation and analysis of tensional tissue. Top left, pipeline used to quantify tissue breakage (red arrow). Fast Random Forest in WEKA plugin in Image J/Fiji was used to segment original images for regions devoid of cells using (red pixels). Bottom and top right, pipeline for segmentation and skeletonization of cell-cell junctions (blue arrows – suppl. Figure 1B). Fluorescent images of EpH4 cells were segmented using machine-learning method (WEKA plugin in Image J/Fiji) to recognize pixels marked by E-Cadherin. Morphological filtering (MorphoLibJ plugin) followed by manual correction were used to rectify remaining segmentation inconsistencies. Segmented images were then skeletonized. B – 37 image features were automatically extracted from segmented and skeletonized images. Random forest model and analysis of features correlation were employed to select the two features that best distinguished untreated (vehicle – low tension) versus nocodazole-treated (high tension) in wild type (Neg-siRNA) samples. These features represent the junction length (J2J length 1) and junction thickness (Junct. thickness 2). A final SVM model was trained using these two features to classify samples as untreated versus nocodazole-treated. All images in A and B have the same size and scaling. Scale bar = 50 µm. C -Importance of each image feature as determined by an ensemble of random forests in the task of distinguishing NegsiRNA images treated with Nocodazole from unstimulated images, during the initial iteration of feature selection. D - Feature correlation analysis was conducted by calculating the correlation for each pair of features. Pairs exhibiting an absolute Pearson correlation greater than T = 0.8 were identified as correlated, forming clusters of inter-correlated features. Within each cluster, only the feature with the highest importance, as determined by the random forest ensemble, was retained. E - Importance of the 12 non-correlated image features as determined by an ensemble of random forests during the second iteration of feature selection. F - Untreated Neg-siRNA samples (vehicle - red circles) and nocodazole-treated samples (blue circles) were projected into a two-dimensional space defined by the most critical features. An SVM was trained to classify the samples, achieving an accuracy of 0.826. The hyperplane that separates the space (solid line) and the soft margin (dotted line) are illustrated. This separation allows for the distinction between images with low(er) versus high(er) tensional state. G – Translation of axes of F gives the distribution of untreated Neg-siRNA samples as Euclidian distance from the classifier (left axis) and provides the distribution of tissues under low tension. Rescaling for the mean value of Euclidean distance is performed to scale values on the more intuitive Apparent tension scale (right axis). Dark and light grey areas indicate the 60% and 90% ranges of confidence, respectively. Scaling and level of confidence can be used to evaluate the tensional state of unstimulated, genetically modified (protein silencing) tissues compared to that measured in unstimulated wild type tissues (Figure 4 and Suppl. Figure 7). H – Plot of low tension (red circles, vehicle) and high tension (blue circles, nocodazole) images as Euclidian distance from classifier in panel F. Untreated and nocodazole-treated Neg siRNA from the same 96-well plates are paired using a solid line. I – Plot of the differences between nocodazole-treated and untreated (vehicle) Neg-siRNA values from panel G provides the distribution of increments scaled according to our classifier (Δ distance classifier, left axis). Rescaling for the mean increment is performed to scale values on a more intuitive Apparent induced tension scale (right axis). Dark and light gray areas indicate 60% and 90% ranges of confidence, respectively. Scaling and level of confidence can be used to evaluate if the silencing of a protein affects the response to increasing tension in the tissue compared to wild type tissue (Figure 6 and Suppl. Figure 8).

The screening was designed to generate three sets of images that allowed pairwise comparison of samples at different conditions (Figure 1B). In each of the 34 plates, the first pair of images (Figure 1B, left) consisted of nocodazole-treated (Nocodazole) and untreated (Vehicle) cells transfected with non-target siRNA (Neg-siRNA). This set of images (N=68, with 34 nocodazole-treated and 34 untreated samples) provided a benchmark control, representing wild-type tissue under both basal and induced tension. These images were used to develop an analytical pipeline to quantitatively characterize tissues under varying tension levels. Each of the thirty-four 96-well plates included this pair of samples (one Neg-siRNA Vehicle and one Neg-siRNA Nocodazole), generating a dataset that served as an internal reference for analysis. The second pair of images (Figure 1B, center) was used to assess the role of each of the cadhesome proteins in regulating tissue formation and tensional state under resting conditions (Figures 3, 4 and Suppl. Figure 7). This was done by comparing wildtype samples that were transfected with non-target siRNA (Neg-siRNA) with those where silencing RNA reduces the expression of a specific cadhesome proteins (GeneX-siRNA). The last pair of samples (Figure 1B, right) was used to evaluate the role of cadhesome proteins in responding to increased tension within the tissue (Figures 5, 6, and suppl. Figure 8). In this pair both samples are silenced for a specific protein and the comparison is between the sample where tension was pharmacologically induced (Nocodazole) with its paired non-treated counterpart (Vehicle).

**Figure 3.**
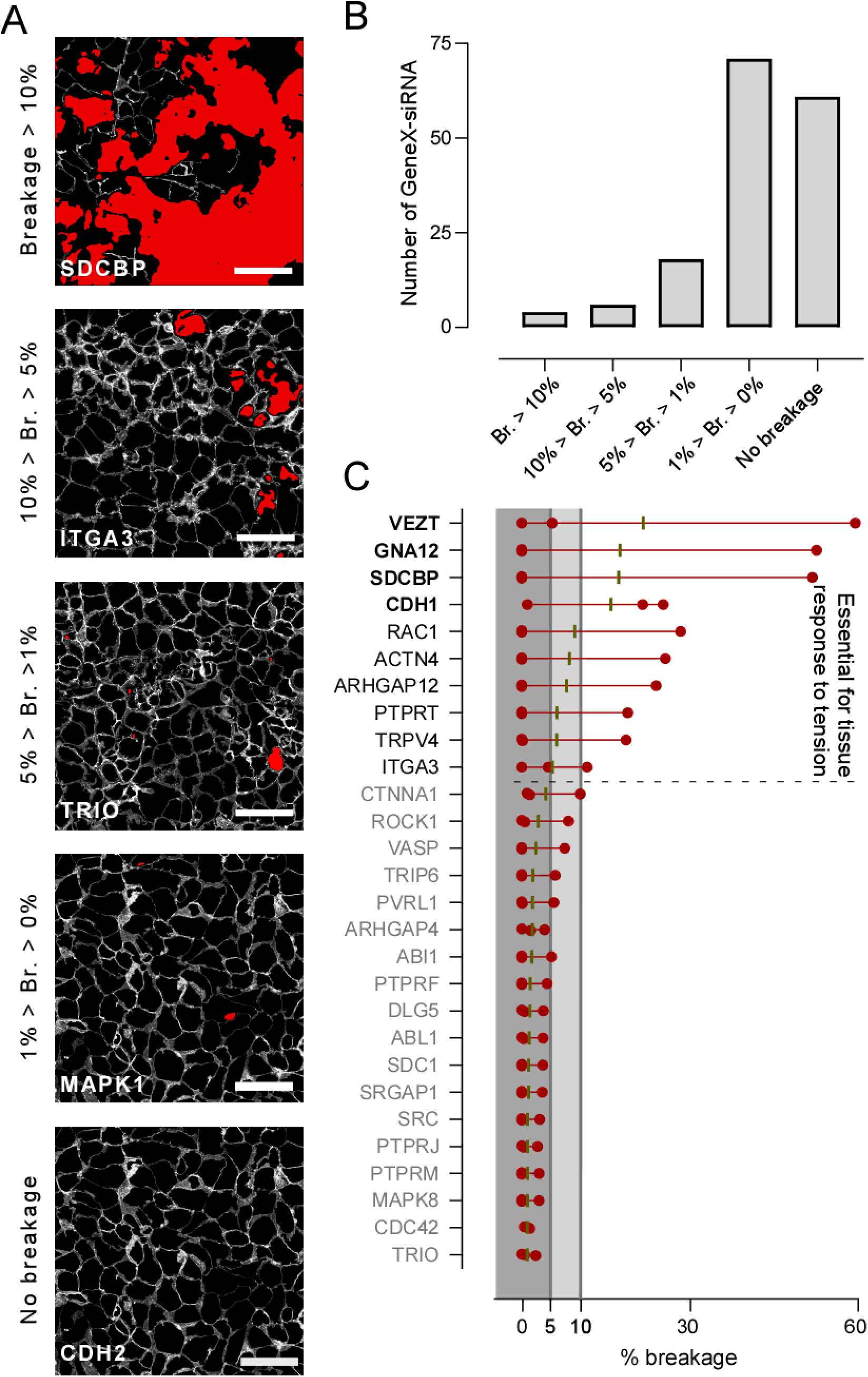
Identification of Proteins Essential for Tissue Formation. A – Exemplary images of EpH4 cells transfected for GeneX-siRNAs which prevent formation of tissues and/or disrupts its integrity. Images are segmented: cell cytosol is in black, cell-cell junctions are in greyscale (linear function of the original intensity) and areas devoid of cells (breakages) are in red. Examples of different percentage of tissue breakages (Br.) are shown from top to bottom (highest to lowest degree), classified as follow: large (Br. > 10 %), medium (10% > Br. > 5%), light (5% > Br. > 1%), low (1% > Br. > 0%), and no breakage (Br. = 0), where Br. is the red area. Scale bar = 50μm. B – Analysis of the number of genes the silencing of which cause different degrees of tissue breakage (as classified in A). C – Analysis of proportion of breakage area caused by gene silencing. Individual measurements (repeats from different experiments, n=3 for each GeneX-siRNA) are depicted by red circles, measurement range is indicated by the connecting red line and the mean of three independent measurements by a black dash. Dark and light grey areas serve as a visual guide for the % of tissue breakage indicated in the x axis of the graph. Protein names (GeneX-siRNA) are in black bold font if the average is above 10%, in black font if between 5 and 10 % and gray font if between 1 and 5%. Genes the siRNA of which causes a breakage in the tissue above 5% are excluded from further analysis using the machine learning classifier (Figure 4) nor for their response to pharmacological stimulation (Figures 5 and 6), as their critical role for formation and maintenance of tissue integrity is already evident.

As result of protein silencing and nocodazole treatments a large spectrum of tissue phenotypes was observed (Figure 1C) when compared with the Neg-siRNA samples. As expected, silencing of key proteins, such as E-Cadherin (CDH1) or nonreceptor tyrosine kinase ABL (ABL1), caused a complete or partial rupture of the tissue, respectively. Tissue ruptures were also observed in some GeneX-siRNA samples when tissue tension was raised by nocodazole stimulation. Besides these extreme phenotypes, proteins silencing and/or nocodazole stimulation caused subtle changes in the organization and architecture of the tissues. For instance, silencing of less crucial proteins, such as JUP, and nocodazole stimulation produced only subtle differences thus requiring the development of an appropriate analytical scheme to quantify differences in tissue tensional state.

### Development of a classification pipeline to evaluate tissue tensional state

To assess the effects of gene silencing and nocodazole stimulation, we developed a multistep analytical pipeline that includes: (1) detection of tissue breakages, which do not occur in normal epithelial tissue; (2) assessment of the tissue’s apparent tensional state relative to internal standards (Neg-siRNA vehicle); and (3) evaluation of changes in apparent tension in response to nocodazole, comparing naïve tissues (Neg-siRNA vehicle vs. Neg-siRNA nocodazole). To quantify tissue breakages, we used *Trainable WEKA Segmentation* ^15^, a machine learning plugin in ImageJ/FIJI (see methods for details on Machine Learning method, features and parameters used), to segment the images for pixels within areas devoid of cells (tissue breakage) and those occupied by cells (Figure 2A, top). Regions not occupied by cells indicated that a specific treatment (silencing or increased tension) caused tissue breakage. The percentage of area of tissue breakage as compared to the total was calculated for all images. If the average percentage of the three repeats was above 5% tissue breakage, the set of images was not further analyzed, as the importance and role of the gene/treatment was already demonstrated beyond any doubts. For the remaining images, we developed a multistep segmentation protocol (Figure 2A, bottom and right and Suppl. Figure 5) and an analytical pipeline to infer the apparent tensional state of the tissue (Figure 2B to I). Images of the nuclei were segmented using simple thresholding method in FIJI/ImageJ. E-Cadherin fluorescence images were segmented using WEKA machine learning (ImageJ/FIJI plugin) trained to differentiate pixels belonging to the cell-cell junction and those of the cytosol (see methods for details). Segmentation was then followed by morphological filtering ^16^ (see methods for details), manual correction for the remaining inconsistencies, and skeletonization. This process generated four images for each sample: a binary image of the nuclei, a binary image of the cell-cell junction, a masked image of the cell-cell junction where the cytosol pixels are set to zero and those of the cell-cell junction retain the original values and, finally, a skeletonized image. Thereafter, we developed an analytical pipeline to classify images based on whether the tissue showed characteristics of a normal, wild-type state (Neg-siRNA vehicle) or was more similar to tissue under tension (Neg-siRNA Nocodazole). This classifier allowed us to quantitatively score the tissue’s tensional state. The features extracted from the three sets of images (nuclei and cadherin segmented and skeletonized) underwent a feature selection process (full list and description can be found in Suppl. Table 2). Initially, features with minimal or no variance across all Neg-siRNA samples (both Vehicle and Nocodazole) were removed, resulting in 37 remaining features. Subsequently, several decision trees models were trained, and the importance of each feature in contributing to the random forest classifier was measured (Figure 2C). The correlation between each pair of features was then calculated (Figure 2D). If two features exhibited an absolute (negative or positive) correlation greater than 0.8, they were marked as correlated. For each group of correlated features, only the feature with the highest importance, as determined by the models, was retained, and the remaining were removed as they do not provide further information. This led to a final set of 12 uncorrelated features which were used to train a new ensemble of 10 independent random forests (Figure 2E). From this analysis the two most important features were selected: J2J Length 1 and Junction Thickness 2 (Figure 2E). These two features represent, in general terms, the average length and thickness of the branches in the segmented image. A linear Support Vector Machine (SVM) ^17^ model was trained using only these two features to classify samples as either Nocodazole or Vehicle, achieving an accuracy of 0.826 (± 0.08) through 10-fold cross-validation. SVM was selected for its robustness to outliers, as it allows for certain points to be misclassified in order to construct a hyperplane that optimally separates the classes and achieves strong generalization. Thus, the trained SVM constructs a hyperplane that separates the feature space into regions corresponding to samples classified as Vehicle (low tension) and Nocodazole (high tension) (Figure 2F). Translation of axes of Figure 2F gives the distribution of untreated Neg-siRNA samples as Euclidian distance from the classifier and provides the distribution of tissues under low tension (Figure 2G, left axis). Rescaling of the mean value of Euclidean distance to 0 was performed to scale values on the more intuitive Apparent tension scale (Figure 2G, right axis). This distribution of these values can be used to determine differences related to the effect of gene silencing, i.e., GeneX-siRNA Vehicle samples (Figure 4 and Suppl. Figure 7). Next, we calculated the pairwise (i.e., of samples on the same 96-well plate) difference between Neg-siRNA Vehicle and Neg-siRNA Nocodazole to measure the average response of tissue to induce tension (Figure 2H). These values determine distribution of the difference between Neg-siRNA Vehicle and Neg-siRNA Nocodazole samples measured on the scale determined by our classifier (Figure 2I, left axis). Rescaling of the mean value of Euclidean distance to 0 was performed to scale values on the more intuitive Apparent induced tension scale (Figure 2I, right axis). These were used to evaluate if the silencing of a protein affects the ability of the tissue to respond to increasing tension as compared to a wild-type tissue (Figure 6 and Suppl. Figure 8).

**Figure 4.**
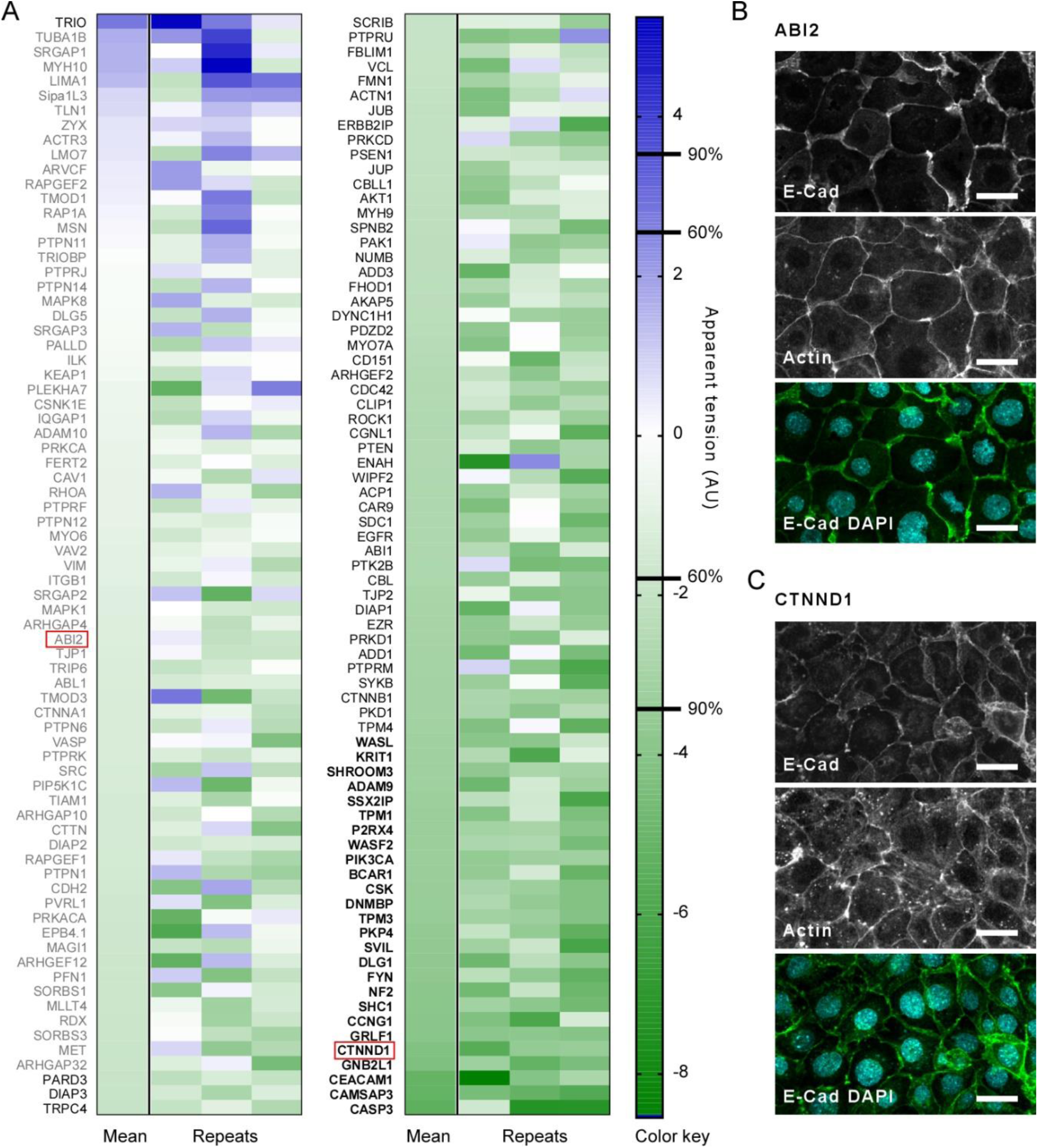
Identification of Proteins Playing a Role in Regulating Tensional State of the Tissue. A – Heatmap reports the apparent tension in tissues with specific proteins silencing calculated as defined by our classification method (Figure 2G). Individual measurements (Repeats) and average of the three independent replicates (Mean) are displayed as arbitrary units (AU) according to the apparent tension scale bar on the right (Color key). Confidence levels from the classifier of 60% and 90% are indicated in the color key bar. Protein names are in black bold font for confidence levels above 90%, in black font for levels between 60 and 90 % and grey fonts for levels below 60%. Only samples whose degree of breakage is below 5% (Figure 3) were analyzed using this classification method. B and C – exemplary images of tissues after protein silencing (highlighted by red boxes in A) used for validation of tissue apparent tension. Imaging of actin cytoskeleton in CTNND1 KD (apparent tension = -4,24; > 90% confidence) revealed the appearance of tension-bearing actin structures (e.g., stress fibers), whereas ABI2 (Apparent tension = -1.28; < 60% confidence) did not. Scale bar = 20 µm.

**Figure 5.**
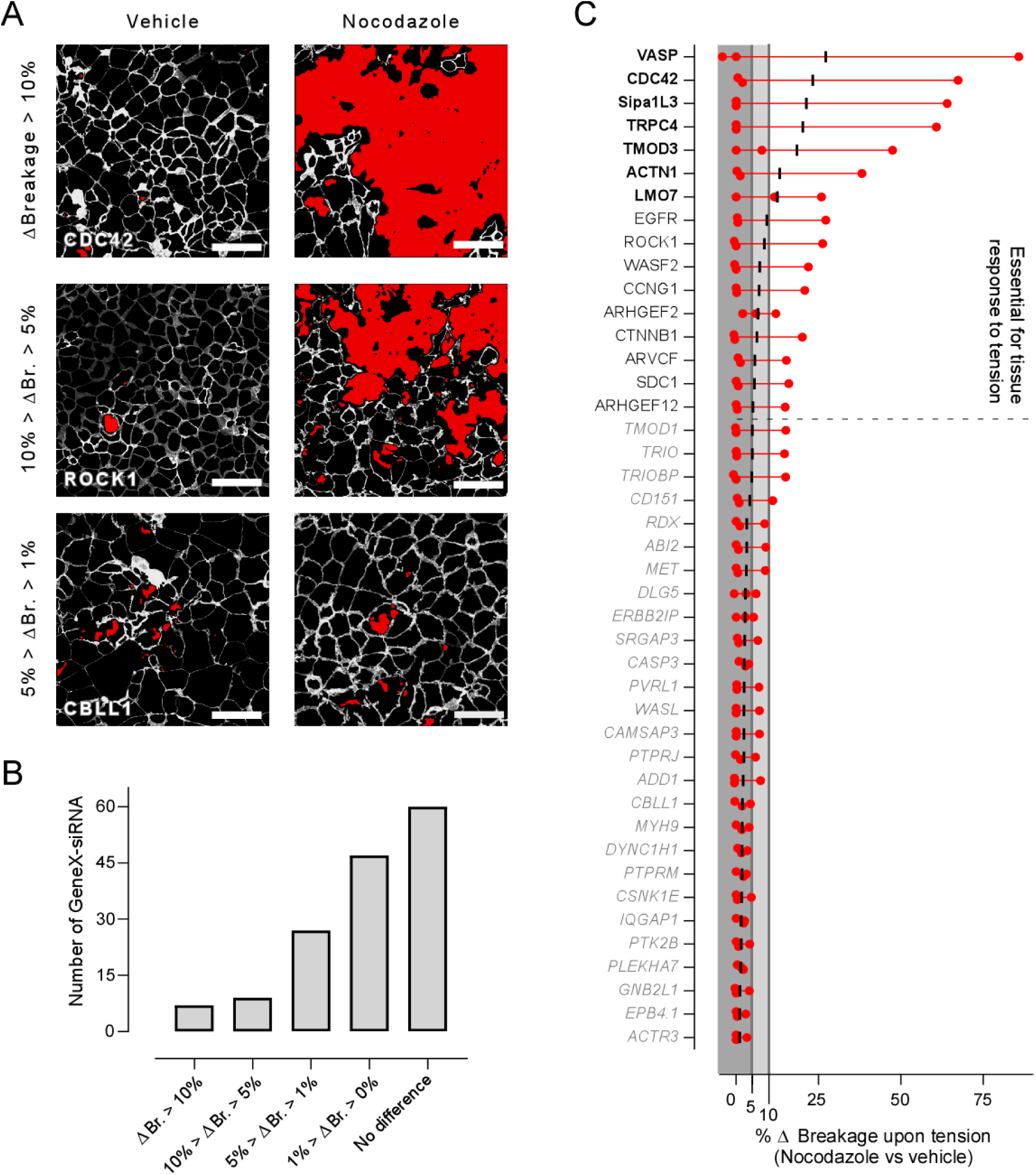
Identification of Proteins Essential for Maintaining Tissue Integrity upon Tension Increase. A – Exemplary images of EpH4 cells transfected for GeneX-siRNAs which led to tissue breakage when tension in the tissue was pharmacologically increased by using nocodazole (2hours 5µM). Paired images of the untreated (vehicle) and nocodazole-treated samples (Nocodazole) are shown on the left and right, respectively. Images are segmented: cytosol is in black, cell-cell junctions are in greyscale (linear function of the original intensity) and areas devoid of cells (breakages) are in red. Examples of different degree of increased (Δ) tissue breakages upon pharmacological stimulations, versus the untreated, are shown, classified as follow: large (ΔBreakage > 10 %), medium (10% > ΔBreakage > 5%), light (5% > ΔBreakage > 1%), low (1% > ΔBreakage > 0%), and no difference (ΔBreakage = 0). Scale bar = 50μm. B – Analysis of the number of proteins the silencing of which cause different degrees of increased tissue breakage upon pharmacological stimulation (as classified in A). C – Analysis of increased broken area for each protein silencing. Individual measurements (repeats from different experiments, n=3 for each GeneX-siRNA) are depicted by red circles with measurement range indicated by the connecting red line and the mean of three independent measurements by a black dash. Dark and light grey areas serve as a visual guide for the % indicated in the x axis of the graph. Protein names are in black bold font if the increase in area devoid of cells is above 10%, in black font if between 5 and 10 % and in grey fonts between 1 and 5%. Relevant increments are defined as those that, on average, are above 1% in the three repeats and severe when above 5% (dashed line separates proteins essential for tissue response to induce tension). In this later case, samples are no further analyzed for tissue appearance upon nocodazole stimulation using the machine learning classifier (Figure 6).

**Figure 6.**
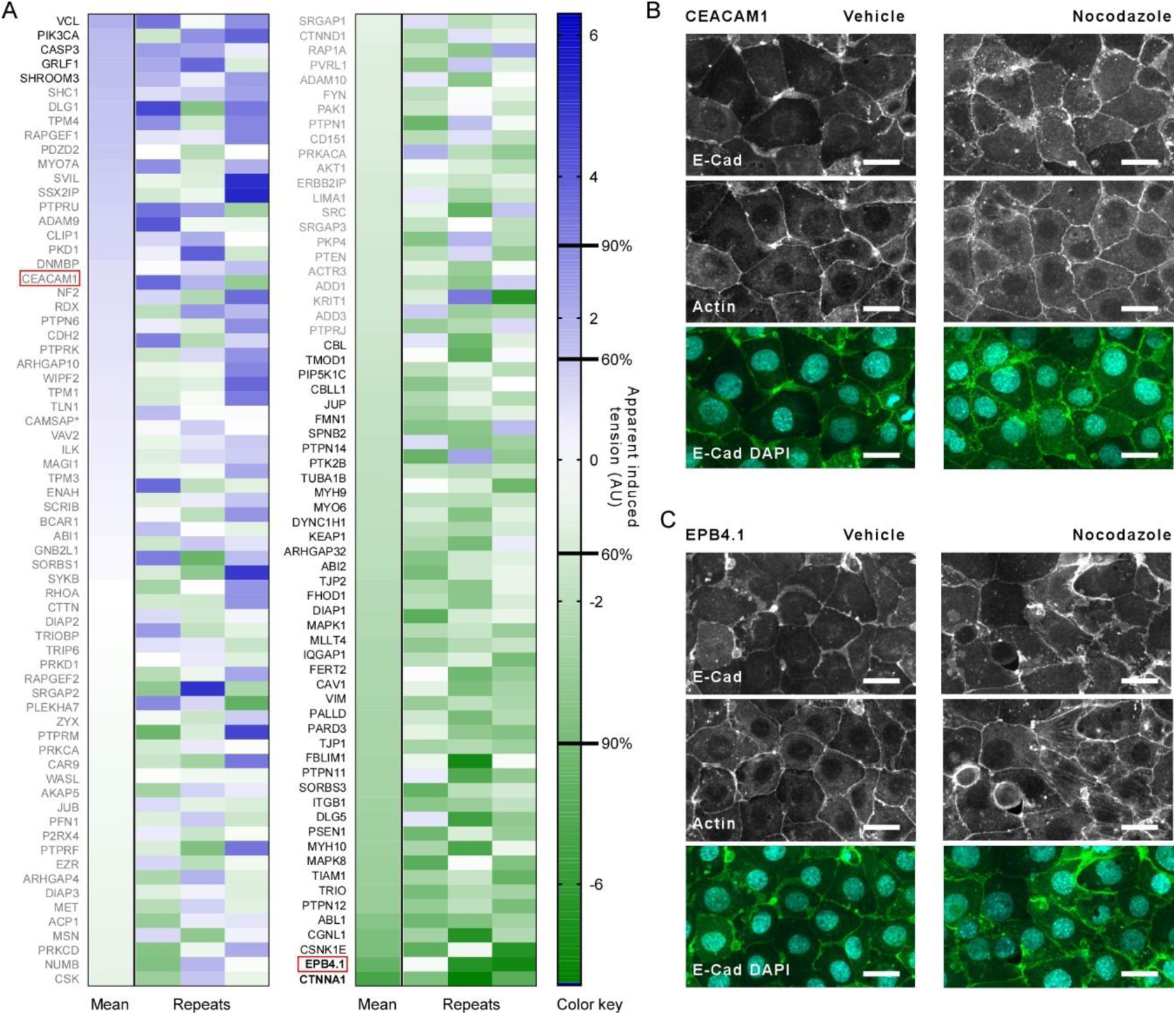
Identification of Proteins Playing a Role in Regulating Response to Tension Increase in the Tissue. Heatmap reports the apparent induced tension in genetically modified tissues (proteins silencing) as a difference between nocodazole treated and untreated samples calculated according to our classification method (Figure 2I). Individual measurements (Samples) and average of the three independent replicates (Mean) are displayed by arbitrary units (AU) according to the apparent induced tension color-coded scale on the right (Color key). Confidence levels of 60% and 90% are displayed in the color key bar. Protein names are in black bold font for confidence level above 90%, in black font for levels between 60 and 90 % and grey fonts for levels below 60%. Only samples whose % of Δbreakage is below 5% are analyzed using this classification method (Figure 5). B and C – exemplary images of protein-silenced tissues under untreated (vehicle) and tension-induced (nocodazole) samples (highlighted by red boxes in A) used for validation of tissue apparent tension. Imaging of actin cytoskeleton in EPB4.1 KD (apparent induced tension = -4,54; > 90% confidence) revealed the appearance of tension-bearing actin structures (e.g., stress fibers) upon nocodazole stimulation, whereas CEACAM1 (Apparent tension = 0.80; < 60% confidence) did not display differences between vehicle and nocodazole samples. Scale bar = 20 µm.

### Identification of essential proteins for tissue formation and regulation of the basal tensional homeostasis of cell monolayer

We applied our analytical pipeline to identify key molecular components involved in the formation and tension regulation of tissue under homeostatic conditions (basal tension state). This was done by comparing the characteristics of GeneX-siRNA Vehicle tissues with those of Neg-siRNA Vehicle tissues (Figure 1B, central pair). First, we quantified the percentage of tissue breakage, defined as areas without cells in the segmented images (red regions in Figure 3A). These breakages were classified as large (above 5% of the area, e.g., SDCBP and ITGA3), moderate (1–5%, e.g., TRIO), mild (0–1%, e.g., MAPK1), or absent (e.g., CDH2). The majority of protein silencing resulted in either mild (71 genes) or no (61 genes) tissue breakages, with only a small fraction leading to moderate (18 genes) or large breakages (Figure 3B). Notably, silencing of VEZT, GNA12, SDCBP, and CDH1 caused breakages larger than 10%, while RAC1, ACTN4, ARHGAP12, PTPRT, TRPV4 and ITGA3 resulted in breakages between 5% and 10% (Figure 3C and Suppl. Table 3). These extreme phenotypes may arise from disruptions to key proteins involved in maintaining cell viability, adhesion to the substrate, and/or formation of stable cell-cell junctions. Further investigation is needed to clarify the exact mechanisms behind these observations. Samples from these proteins, which are critical for tissue formation, were excluded from further analysis by the machine learning classifier and from nocodazole stimulation tests as their crucial role has already been established, and the absence of large tissue portions could lead to computational artifacts.

Next, we used our machine learning classifier (Figure 2G) to analyze the appearance of tissues under homeostatic conditions (i.e., without induction of tension) that did not show a breakage larger than 5%. This analytical scheme assesses the statistical confidence in whether the observed tissue resembles the untreated baseline condition (Neg-siRNA Vehicle) or the elevated-tension condition induced by nocodazole treatment (Neg-siRNA Nocodazole). By comparing the tissue characteristics to these two reference states, the analysis determines how closely the sample aligns with either the relaxed or the high-tension profile, providing a quantitative measure of similarity to each condition. Of the 150 genes silenced, 78 showed a decrease in Apparent Tension of which 52 with confidence above 60% and 26 above 90% (Figure 4A, Suppl. Table 3 and Suppl. Figure 7). In contrast only one gene (TRIO) showed an increase in apparent tension (> 60% confidence).

To validate our analytical scheme, we assessed tissue organization and actin cytoskeleton structure in selected samples, comparing them to the Neg-siRNA Vehicle condition (Supplementary Figure 3). Silencing ABI2, a gene found to have no impact on tension (confidence level below 60%), showed no changes relative to Neg-siRNA Vehicle (Figure 4B). In contrast, silencing CTNND1, a gene that associated with increased tension (confidence level above 90%), resulted in distinct alterations in both tissue organization and actin structure compared to Neg-siRNA Vehicle, confirming its impact on apparent tissue tension (Figure 4C).

### Identification of proteins regulating response to induced tension in the tissue

In addition to investigating tissue formation and tension regulation under homeostatic conditions, our screening aimed to explore the role of cadhesome proteins in regulating the tissue’s response to increased tension (GeneX-siRNA Vehicle versus GeneX-siRNA Nocodazole, right pair in Figure 1B). To induce this response, we chemically stimulated cell contractility using nocodazole, which increases the normal pulling forces at cell-cell junctions (Suppl. Figure 3). First, we assessed whether silencing specific genes led to changes in tissue breakage (ΔBr.) following nocodazole stimulation compared to the homeostatic condition (Figure 5). In Figure 5A, images display large increase in tissue breakage in ROCK1 and CDC42 KD samples (above 5 and 10%, respectively), while CBLL1 KD showed moderate increase (ΔBr. between 1 and 5%). Overall, most protein silencing resulted in mild or no increase in tissue breakage (47 genes with ΔBr. between 0% and 1%, and 60 genes with no increase). However, 27 genes showed moderate increases (1% to 5%), and 16 genes had large increases. In this latter set, 9 (EGFR, ROCK1, WASF2, CCNG1, ARHGEF2, CTNNB1, ARVCF, SDC1 and ARHGEF12) ranged between 5% and 10%, and 7 (VASP, CDC42, Sipa1L3, TRPC4, TMOD3, ACTN1 and LMO7) ranged above 10% (Figure 5B). The cause for this extreme disruption could indicate that these proteins play a crucial role in mediating cell adhesion to the substrate and/or formation of stable cell-cell junctions when cells are subjected to increased tension. Samples from these proteins, which are critical for tissue response to tension, were excluded from further estimation of changes in response to tissue induction using the machine learning classifier.

Next, we analyzed whether the response to Nocodazole-induce tension by genetically modified tissue (GeneX-siRNA Nocodazole versus GeneX-siRNA Vehicle) was different from that of a normal tissue (Neg-siRNA Nocodazole versus Neg-siRNA Vehicle). This was assessed by quantifying and statistically evaluating the variation in apparent tension in the Neg-siRNA Nocodazole and Neg-siRNA Vehicle pairs (Figure 2I). This variation was defined as Apparent Induced Tension. Based on this analysis, we determined that of the remaining 134 genes, which exclude genes whose silencing resulted in rupture above 5% in homeostasis or increase rupture upon Nocodazole stimulation, 5 genes showed an increased response to Nocodazole (higher Apparent Induced Tension) with confidence level of 60% (Figure 5A, Suppl. Table 3 and Suppl. Figure 8). On the other hand, 43 and 2 (EPB4.1 and CTNNA1) genes showed a decreased response to Nocodazole (lower Apparent Induced Tension) with confidence level of 60% and 90%, respectively. This may indicate that those genes showing higher Apparent Induced Tension as compared to the response of a normal tissue could be involved in negative regulation of response to tension. Conversely, those displaying lower Apparent Induced Tension could be involved in positive regulation of response to tension. We validated our assay results by imaging the actin cytoskeleton in randomly selected samples. Furthermore, we validated our analytical scheme by examining tissue organization and actin cytoskeleton structure in selected samples, comparing them to the Neg-siRNA Vehicle and Neg-siRNA Nocodazole conditions (Supplementary Figure 3). EPB4.1 and CEACAM1 were chosen to represent an altered response to tension (confidence level above 90%) and a non-altered response (confidence level below 60%), respectively (Figure 4B). Silencing CEACAM1 produced a response to nocodazole similar to that of Neg-siRNA (Supplementary Figure 3). In contrast, E-Cadherin and actin staining in EPB4.1-deficient tissues showed significant changes in tissue organization and actin structure.

### Network representation and bioinformatic analysis

As a result of this large screen, we identified several genes involved in the mechanical regulation of tissue organization and response to increased tension (Suppl. Table 3). Gene silencing yielded a broad spectrum of outcomes, prompting us to classify the genes by their relative importance based on the magnitude of effects and the confidence level provided by our analytical tool. First, we grouped genes whose impact is unequivocal; silencing these genes caused tissue breakages above 5% either under homeostatic or induced tension conditions. This category, labeled "Tissue Breakage," includes genes crucial for tissue integrity. A second group, "ML Confidence > 90%," contains genes that show changes in apparent tissue tension or induced tension determined with a high confidence level (above 90%). Importantly, those thresholds for determining positive hits were set to high levels to reduce the risk of selecting false positives. Lastly, we included a class called "At Least Two Positive Hits" for genes with effects supported by two different criteria, albeit at lower confidence levels. Here, criteria include mild tissue breakages (1–5%) under homeostasis or Nocodazole treatment, and alterations in apparent tension or apparent induced tension determined with a lower confidence (60–90%). Additionally, we used UniProt and String databases to categorize genes by function (Suppl. Table 3) and to map the predicted physical interactions (Suppl. Table 4). Functional categories include interfacial complex proteins, cytoskeletal proteins, signaling proteins, and a miscellaneous group (others) for proteins such as ion channels and proteases. Physical interactions were scored based on the probability score provided by the String. Finally, we used TIMER to examine the impact of specific gene silencing on breast cancer patient survival outcomes (Suppl. Table 5). Based on the results of the screening and bioinformatic information, we constructed a network where nodes represent genes whose silencing yielded positive hits, and edges represent their physical interactions (Figure 7). The results of our screening are organized in three subnetworks, as identified based on the high STRING scores of physical connections and the high degree of connectivity of key nodes: the first is centered around the mechanotransduction module formed by E-cadherin and catenins; the second, around the signaling network of EGFR and CBL; and the third, involving the signaling network including RAC1 and CDC42.

**Figure 7.**
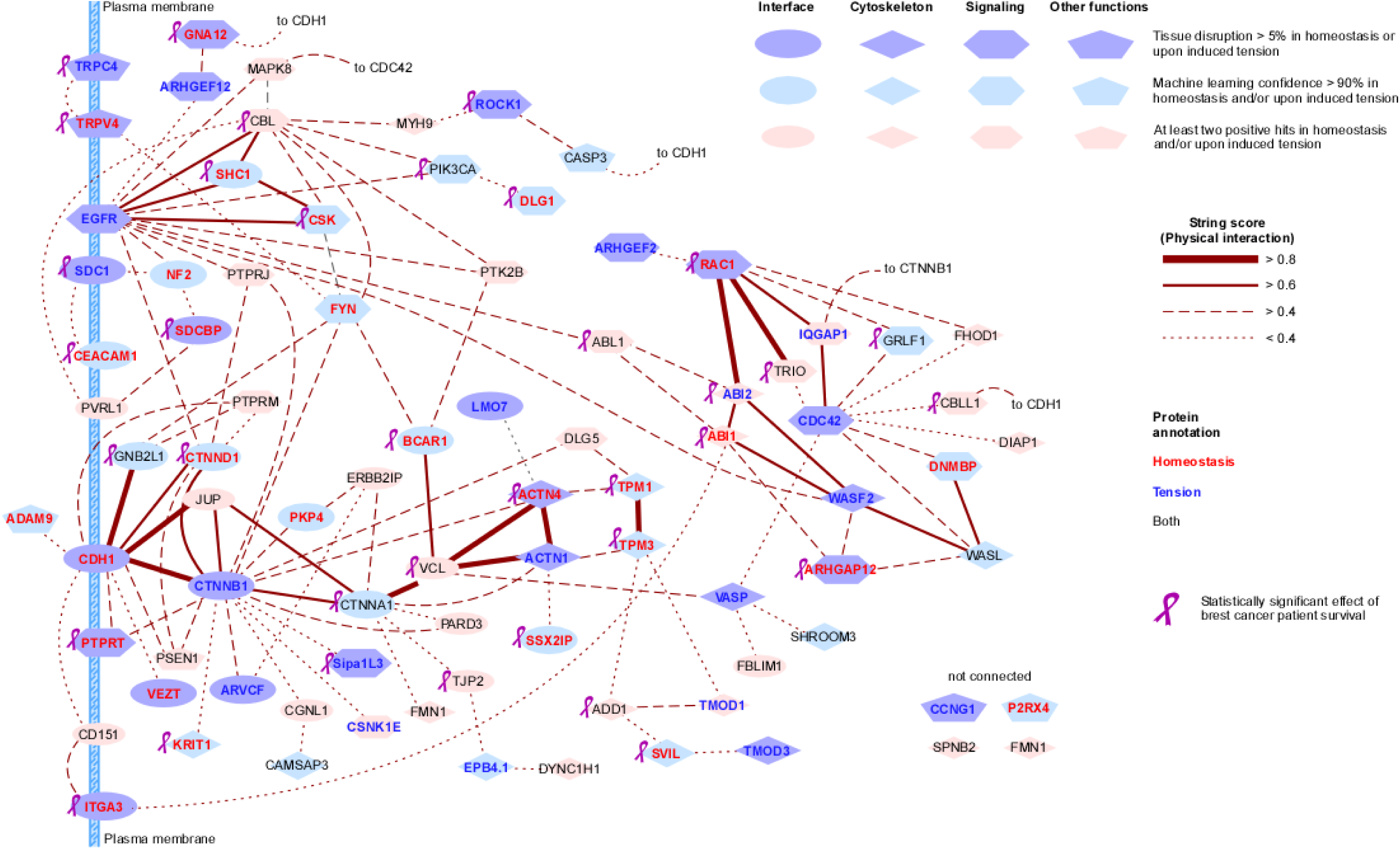
Network Representation of Cadhesome Protein Positive to our Screening. Proteins positive to our screening (criteria indicated in the Figure) are shown as nodes in a network of closest physical interactions. Different shapes of the nodes indicate the proteińs best known function (Suppl. Table 3) categorized as interface (i.e., interfacial complex), signaling, cytoskeleton and other functions. Network was built using STRING (https://string-db.org/). Only physical interactions (experiments) and no confidence threshold (score) were used (Suppl. Table 4). Top three interactions defined by their score are shown (> 0.4). If at least two interaction whose score is > 0.4 are not found, one weak interaction (< 0.4) is shown. Thickness of the edges represents the strength of the interaction as indicated in the figure (String score). Names of protein whose silencing produced an effect on untreated tissues, i.e., tissue formation and homeostasis (Figure 3 and 4), are displayed in red and those affecting the response to pharmacologically-induce tension (Figure 5 and 6) in blue. Those that show positive hits in both assays are displayed in black. The pink ribbon indicates genes whose silencing significantly affects patient survival, as determined by Kaplan–Meier analysis in TIMER (Suppl. Table 5) ^20^.

### Mechanotransduction module

The mechanotransduction module essential to the structure and function of the interfacial complex is centered around the E-cadherin receptor (CDH1) and the adaptor proteins β-catenin (CTNNB1), α-catenin (CTNNA1), and vinculin (VCL). As expected, silencing CDH1 caused large disruptions in tissue integrity under homeostatic conditions ^18^, whereas silencing CTNNB1 impacted tissue integrity only when tension was induced. This is consistent with our recent findings that β-catenin can participate in mechanotransduction independently of α-catenin ^11^. Silencing CTNNA1, a well-established mechanotransducer ^19^, altered apparent tension in homeostasis and response to induced tension with a high degree of confidence. Interestingly, silencing VCL led to decreased tissue tension in homeostasis, but an increase under induced tension. Although these effects were detected with lower confidence (below 60%), this behavior reflects the complex role of vinculin in regulating tissue dynamics across the major mechanical interfaces of the cell (i.e., focal adhesions and adherens junctions). Notably, reduced expression of CTNNA1 and VCL, but not of CDH1 and CTNNB1, impacted significantly cancer patient survival as indicated by the analysis based on Kaplan–Meier survival of cancer patient obtained from TIMER ^20^ (Suppl. Table 5). The lack of correlation between CDH1 and CTNNB1 with cancer patient survival could be due to biopsies that predominantly sample late stages of cancer progression, where typical epithelial markers are largely absent. However, cohorts have also shown that mutations in CDH1 occur more frequently in older patients, while young-onset patients have a higher proportion of CTNNB1 mutations, thus a correlation with survival could be lost if clinical outcomes are not adjusted for patient age ^21^.

Proteins of the mechanotransduction module are central to a complex subnetwork of proteins positive for our analytical pipeline. This network predominantly includes key components of the interfacial complex, such as the catenin proteins (ARVCF, CTNND1, and JUP), as well as adaptor proteins (VETZ, GNB2L1, BCAR1) that mediate interactions with other cellular structures. Additionally, it encompasses an integrin (ITGA3), a signaling protein (Sipa1L3), and a cytoskeletal component (KRIT1). ARVCF (Armadillo Repeat gene deleted in Velo-Cardio-Facial syndrome) silencing resulted in large tissue breakages upon nocodazole stimulation, demonstrating its role in regulating tension at cell-cell junctions during morphogenesis ^22^, as also indicated by its involvement in developmental disorders, such as cleft palate and facial dysmorphology ^23^. CTNND1 (encoding p120 catenin) was found to regulate tensional homeostasis, likely by preventing cadherin endocytosis and recycling ^24^ and by connecting AJ to signaling proteins (e.g., EGFR, PTPRJ, PTPRM). Both ARVCF and p120 interact with small GTPases RhoA and RAC, affecting actin dynamics during development ^25^. JUP (encoding junction plakoglobin), a β-catenin homologue, interacts with E-cadherin and α-catenin and it can be part of both adherens junctions and desmosomes ^26^. Vezatin (VETZ), an adherens junction transmembrane protein, was essential for maintaining tissue tensional homeostasis, possibly by mediating actomyosin contractility-cadherin interactions ^27,28^. GNB2L1 (RACK1, receptor for activated C kinase 1) played a role in supporting tissue tension regulation under both homeostatic and induced tension conditions. BCAR1 (a.k.a. p130Cas/BCAR1), part of the Cas family, was shown to be critical for tensional homeostasis in our study. This scaffold protein impacts multiple pathways, including cell motility, apoptosis, and cell cycle control^29^, and its dysregulation affects pathways linked to cardiac development, liver function, and cancer (Barrett et al., 2013. Additionally, integrin ITGA3 (via CD151) was associated with mechanotransduction module, with silencing causing large tissue breakages under homeostatic conditions. The signaling protein Sipa1L3 induced tissue breakage under increased tension, likely via Rap1-GTPase ^30,31^. KRIT1 was observed to be crucial for maintaining tissue tensional homeostasis, possibly by interacting with integrins (ICAP1α) and microtubules ^32^. Finally, differential expression of ITGA3, CTNND1, VETZ, Sipa1L3, BCAR1, KRIT1, and TJP2 impacts breast cancer patient survival.

Within the mechanotransduction module, vinculin and, to a lesser extent, α- and β-catenin connect to the actin cytoskeleton via actin-crosslinking proteins α-actinin 1 and 4 (ACTN1 and ACTN4). These proteins, in turn, link to the actin-binding tropomodulin 3 and tropomodulin 1 (TPM3 and TPM1), respectively ^33,34^. Silencing of both actinins resulted in tissue breakages with ACTN4 affecting tissue homeostasis and ACTN1 response to included tension. On the other hand, both tropomodulins affected tissue tensional state in homeostasis. Notably, expression changes in both tropomodulins, as well as ACTN4, are associated with breast cancer patient survival. In addition to this circuit, α- actinins also interact with LIM domain only 7 (LMO7) and the cancer-associated SSX2-interacting protein (SSX2IP) ^35,36^. Both proteins are known or predicted to associate with adherens junctions and nectin-afadin adhesion complexes, playing a role in organizing the actin cytoskeleton at cell-cell adhesion sites)^35,36^. Tropomodulin 3 connects to another small network formed by the adducin 1 (ADD1), supervillin (SVIL) and the two tropomodulin 1 and 3 (TMOD1 and TMOD3), with TMOD3 showing the larges effect in our screening and causing tissue breakages in response to induced tension. Changes in expression of ADD1 and SVIL are related to changes in cancer patient survival ^37,38^. Interestingly, ADD1 phosphorylation has been demonstrated to be part of the lamellipodia formation and migration of breast cancer cells in response to EGF signaling ^39^ .

### EGFR signaling module

A second subnetwork is centered around epidermal growth factor receptor (EGFR), E3 ubiquitin-protein ligase CBL (CBL), SHC-transforming protein 1 (SHC1), c-terminal src kinase (CSK) and it is connected with the mechanotransduction module principally through the tyrosine-protein kinase Fyn (FYN), a Src family tyrosine kinase and, to a lesser extent, the G protein subunit alpha 12 (GNA12). EGFR is a well-known regulator of epithelial tissue physiology. Upon activation by its ligands, primarily Epidermal Growth Factor (EGF), EGFR initiates signal transduction pathways that promote cell proliferation, differentiation, migration, and survival) ^40^. Silencing of EGFR caused breakages in the tissues only upon induction of tension within the tissue. Closely connected to EGFR we found CSK, SHC1 and CBL that are central mediators in tyrosine kinase signaling. CSK is involved in several processes, including adherens junction maintenance and positive regulation of hippo signaling ^41^. SHC1 acts as an adaptor protein that links activated receptor tyrosine kinases (including EGFR) to downstream signaling cascades such as the Ras-MAPK pathway ^42^. Finally, the E3 ubiquitin ligase CBL regulates E-cadherin complex turnover and degradation ^43^). Silencing of CSK and SHC1 were found to alter the tensional state of the tissue in homeostasis, while CBL seemed to have a less pronounced effect on the tissue. The G proteins subunit GNA12 and the Rho guanine nucleotide exchange factor ARHGEF12 are also critical nodes of this subnetwork, and their silencing caused tissue breakages in homeostasis and upon induced tension, respectively. These proteins are known to activate RhoA/ROCK1 pathway, thus linking mechanical signals to the cytoskeletal reorganization necessary for tumor cell invasion. Besides the genes already discussed above, our screening revealed that a number of additional genes were associated with tissue breakage and changes in tissue tension. In homeostasis, DLG1, NF2, CEACAM1, and GNA12 were found to lead to tissue breakage. Upon the induction of tissue tension, ROCK1, SDC1, and ARHGEF12 also resulted in tissue breakage. Under nocodazole treatment, SHC1 was identified as key genes influencing changes in tissue tension with high confidence. Additionally, silencing FYN led to significant changes in the tissue’s tensional state in homeostasis. PIK3CA had similar effect in both homeostasis and nocodazole stimulation. Importantly, database analysis correlated the expression level of several genes within this subnetwork (SHC1, CBL, CSK, ROCK1, PIK3CA, DLG1, SDC1, CEACAM1, SDCBP, GNA12) with survival of breast cancer patients. FYN, which is a node with one of the highest degrees of connectivity, connects to two members of the family of Transient receptor potential (TRP) calcium channels: the canonical TRPC4 and member of the subfamily V TRPV4. This second is known to be activated by several physical cues including temperature and mechanical stresses. Importantly, silencing of both genes resulted in severe disruption in tissue integrity in homeostasis (TRPV4) or upon induction of tension (TRPC4). Analysis of patient survival using TIMER confirmed that both genes play critical roles in a variety of cancer related processes^44^. As mentioned above, increased expression of EGFR upon the surface of tumor cells and the presence of activating mutations of EGFR are both correlated with poor breast cancer patient survival. This is often contributed to a recruitment of FYN and other kinases within the EGFR scaffold which then feed into the MAPK pathway causing progression of the cell cycle and thus promoting proliferation and survival. Although requiring further validation in a uniquely cancer cell model, the results presented herein further suggest that the EGFR-EGF pathway provides signals that help prevent tissue breakage and may enhanced adaptation to the changing tensions present in a fast proliferating tumor, with the result of cancer cell survival.

### Small GTPases signaling module

The signaling subnetwork centered around CDC42 and RAC1 connects to the mechanotransduction module primarily through VASP and to the EGFR subnetwork via ABL1, positioning VASP and ABL1 as integrators of actin-regulatory functions from CDC42, RAC1, and associated proteins such as ARHGEF2, GRLF1 and WASF2. VASP (Vasodilator-Stimulated Phosphoprotein), a member of the Ena/VASP family, promotes actin filament elongation at adhesion sites by interacting with actin-binding proteins like vinculin, which also determines the tension-dependent localization of VASP at focal adhesions or adherens. In our assay, silencing of VASP caused tissue breakages when tension was increased using nocodazole. Similarly, silencing CDC42 also led to tissue breakages upon nocodazole stimulation. CDC42 is a GTPase that regulates actin polymerization via activation of N-WASP and ARP2/3 complex to promote branched actin polymerization, supporting migration, adhesion, and mechanotransduction ^45^. ABL1, on the other hand, links this subnetwork to the EGFR signaling network. As a non-receptor tyrosine kinase, ABL1 is activated downstream of EGFR signaling through the Ras-Raf-MEK-ERK pathway or directly via EGFR-dependent activation of Src family kinases ^46^. Once activated, ABL1 can phosphorylate multiple actin-regulatory proteins, including ABI1 and ABI2 ^47^. Phosphorylation of ABI1 promotes its interaction with TRIO, stimulating RhoA GEF activity ^48^, and activates the WASF2 complex to initiate ARP2/3-mediated actin filament branching and polymerization^49,50^. ABI2 phosphorylation contributes to RAC1 interaction, supporting actin polymerization and regulation of cell adhesion, polarization, and migration^51^. ABL1 also phosphorylates adaptor proteins such as vinculin and β-catenin, underscoring its role as a hub that connects signaling pathways with cytoskeletal modulation and mechanotransduction ^3,52^. Within this subnetwork, silencing of ARHGAP12 and RAC1 was associated with tissue breakages under homeostatic conditions, while CDC42, ARHGEF2, and WASF2 caused breakages in response to increased tension. Silencing DNMBP increased tissue tension in homeostasis alone, whereas GRLF1 and WASL knockdown altered tension in both homeostasis and during nocodazole stimulation. Interestingly, TRIO silencing was found to increase tissue tension under homeostatic conditions. Although this finding was determined with low confidence, it suggests a role for TRIO as a “shock absorber,” potentially mitigating excessive tissue tension. In epithelial cells, TRIO has been shown to mediate barrier destabilization via RAC1-mediated repression of E-Cadherin expression, further highlighting its role in tissue mechanics and barrier function ^53,54^. Finally, lower levels of expression in patient biopsies of ABL1, ABI1, ABI2, RAC1, ARHGAP12, CBLL1, GRLF1 and TRIO impacts breast cancer patient survival.

Overall, our work provides insights into the function of proteins of the cadhesome network on regulating the mechanics of an epithelial tissue model. Furthermore, it provides the blueprint for how to design, conduct and analyze the complex relation between gene expression, protein function and biomedical significance of a large and complex network of proteins using a high-throughput platform and how to use artificial intelligence as an analytical tool capable of quantitatively accessing protein function.

## Discussion

This study provides a comprehensive examination of the proteins within the cadhesome network and their role in regulating epithelial tissue mechanics, offering novel insights into the function of mechanotransductive proteins and their broader biomedical relevance. Through a systematic analysis, we identified three primary subnetworks that are central to the regulation of tissue integrity and mechanical behavior: the mechanotransduction module, the EGFR signaling network, and the network of small GTPases that regulate the actin cytoskeleton. These networks highlight the complexity of the force-transduction machinery, illustrating how mechanical processes are integrated with biochemical signaling cascades. This integration is crucial for the mechanical response of epithelial tissues, enabling them to regulate tensional homeostasis and adapt to physiological changes in tension during dynamic process involving coordinated cell migration, such as morphogenesis, wound healing, and tissue regeneration^55–58^. Similarly, these results also highlight how modifications in the expression patterns of proteins involved in the regulation of tissue mechanics can have severe implications for cancer progression, ultimately impacting patient prognosis.

Given the complexity of the networks involved in tissue mechanics, our study establishes a foundational blueprint for designing and executing high-throughput investigations of other complex cellular functions by exploring the relationship between gene expression, protein function, and functional outcomes – an emerging, novel approach known as functional omics ^10,59,60^. By leveraging advanced technologies such as high-throughput imaging, artificial intelligence, image analysis tools, bioinformatics, and functional assays, our work not only deepens our understanding of mechanotransduction and its pivotal role in both homeostasis and disease, but also offers valuable insights that could drive the identification of novel biomarkers and therapeutic targets.

## Materials and Methods

### Cell culture and sample preparation

EpH4 cell line is a non-tumorigenic cell line derived from spontaneously immortalized mouse mammary gland epithelial^61^. Eph4 mouse mammary epithelial cell line is a kind gift from Jean-Paul Thiery (IMCB, A-STAR, Singapore). Eph4 cells were cultured in DMEM with 10% FBS and 1% Penicillin-streptomycin (Life Technologies). No cell lines used in this study were found in the database of commonly misidentified cell lines maintained by ICLAC and NCBI Biosample. The cell lines were not authenticated. The cell lines used were regularly tested for mycoplasma contamination by PCR methods.

### RNA sequencing

R RNA was extracted from 10 million EpH4 cells using a QIAGEN RNAesy Mini Kit following manufacturer instructions. For RNA-seq Quantification we followed Sample Requirements for BGI-SEQ 500 service. Integrated total RNA samples were treated by DNase to avoid protein contamination during RNA isolation. The samples collected had total RNA ≥ 200ng at concentration ≥ 20 ng/µl and measured purity of OD260/280 ≥ 1.8, OD260/230 ≥ 1.8. Samples were then shipped to BGI Tech Solutions Co. Limited (HongKong) on dry ice for RNA-seq quantification.

### siRNA library and cellś transfection

EpH4 cells were transfected in suspension at early passage (below P22). 50 µl transfection mixture (5 µl siRNA [1 µM] in 20 µl OPTIMEM with 0.5 µl Lipofectamine [Invitrogen] in 24.5 µl OPTIMEM) were added to 50 µl of cell suspension (18000-20000 cells). The final siRNA concentration was 50 nM Cells were incubated at 37 degrees at 5% CO2. After 6h OPTIMEM was replaced with fresh antibiotic-free DMEM containing 10%FBS. For the transfection, we used a mouse siRNA library targeting 176 proteins previously identified as the cadhesome^8^ with predicted roles in cell-cell adhesion. SMARTpools siRNAs obtained by Thermo Fischer Scientific consisted of mixtures of four different siRNA sequences targeting the individual genes of the cadhesome (GeneX-siRNA). Each 96-well plate, aside from cells transfected with SMART pools siRNAs targeting the cadhesome genes, also allocated controls: samples transfected with non-target siRNA (Thermo Fisher Scientific, here indicated as Neg-siRNA), which served as control for non-specific effects related to siRNA delivery to provide a baseline for target gene silencing, and two samples transfected with the transfection indicators, siTOX Transfection Control, and with siGLO RISC-Free Control siRNA, respectively, as control for transfection efficiency. Cell successfully transfected with siTOX undergo apoptosis and cell death within 24hours (Suppl. Figure 2A), the degree of cell death can then be correlated to transfection efficiency. About siGLO, while the exact mechanism is not exactly known, following complex formation of lipid-based transfection reagents and siRNA, small nanoparticles are formed and are taken up by a receptor-mediated endocytosis. The observed siRNA spots, likely endosomes, (Suppl. Figure 2A) within the interior of the cell suggest excellent transfection efficiency (>90% have at least one or 2 spots, if not more). Twenty-four hours after transfection, if siTOX and siGLO demonstrated that the transfection had been successful, cells were treated and subsequently fixed and stained for E-Cadherin and the nuclei.

### Pharmacological Treatment for induction of tension

24 hours after transfection, and prior to fixation, cells were treated for 2 hours with 5µM nocodazole (M1404, Sigma) to induce tension at the junction increasing the intracellular contractility (Suppl. Figure 1). Nocodazole is a microtubule-depolymerizing agent that induces Rho-dependent actin stress fiber formation and contractile cell morphology. RhoA GTPase plays a vital role in assembly of contractile actin-myosin filaments (stress fibers) and of associated focal adhesion complexes of adherent monolayer cells in culture. GEF-H1 is a microtubule-associated guanine nucleotide exchange factor. Thus, depolymerization of microtubules results in the release and activation of GEF-H1, which in turn activates RhoA (2). We have previously demonstrated by laser nano-scissor experiments that treatment with nocodazole leads to an increase in junctional tension in epithelial monolayers^3^. Here we have validated the effectivity of nocodazole treatment by staining with F-actin and Myosin IIA and imaging of EpH4 wt non-treated (vehicle) and Nocodazole treated (Suppl. Figure 3) and imaging at different focal plane to highlight stress fibers (Basal), junctional actin and myosin organization (Apical).

Additionally, we used mechanical induction of force, by application of tension using an in-house build uni-directional strain device (i.e. Suppl. Figure 4). Cells were grown on an elastic, optically clear, inert silicone membrane, which was fixed in a clamping device. After 1hour of stretching cells were fixed and submitted to staining.

### Immunostaining

After drug treatment, or stretching, cells were fixed in 4% PFA for 15 min, and permeabilized with 0.2% Triton for 3 min at RT. Blocked with 1% BSA for 1 hour at RT and stained for Ecadherin and nuclei (DAPI). Every washing step was performed with PBS with Ca2+ to preserve the Ecadherin homofilic binding between neighboring cells of the tissue.

The antibodies used are the following: Primary Antibody Rat anti-Ecadherin, clone DECMA-1 (MABT26, Sigma Aldrich), Primary antibody Rabbit Anti Myosin IIA (M8064, Sigma Aldrich) Phalloidin Alexa 647 (Thermofisher Scientific), Secondary Antibody Goat anti Rat Alexa 568, anti-Rabbit 488 (Invitrogen), and DAPI (Invitrogen).

### Image acquisition

Images were acquired using a CSU spinning disk (Yokogawa®) confocal head mounted on an inverted microscope (Eclipse Ti, Nikon, Japan). Images were acquired using a Plain Apo Vc 60X oil objective, N.A. 1.40, with different excitation wavelengths for DAPI (405nm), Alexa 488, Alexa 568 and Alexa 647.

High-content image acquisition of fixed cells in 96-well plates was performed using a Metamorph based high-content screening (HCS) module. Nine adjacent z-stacks (step 0.5um, total height 3.5um) have been averaged to a projection and stitched in a 3x3 montage to obtain a 1489*1489px field of view.

### Image segmentation and skeletonization

Trainable WEKA Segmentation plugin, a Random forest based machine-learning plugin, on ImageJ/Fiji^15^ has been employed for cell-cell adhesion segmentation in the images obtained by the previous high-content image acquisition of the 96-well plates. The pipeline consisted of obtaining three different trainings based on the image brightness level (faint, medium, bright). All images were segmented via batch processing on ImageJ, yielding three segmentations per original image, with a fourth result obtained via an AND operation of the previous three; the result with the best performance was then converted to 8-bit binary and submitted to morphological filters with MorpholibJ plugin^16^ (closing operation, octagon element, and radius 4; directional filter with a max type, opening operation, line 26, and direction 32). Manual correction was performed on the images to eliminate any residual noise left during the segmentation process. For the nuclei, DAPI images were segmented using a simple thresholding method in ImageJ/FIJI. For E-Cadherin-stained images, we used WEKA Segmentation to (1) segment images for tissue breakages and (2) for cell-cell junction. In the first case, training was done by segmenting images into two classes: cell-occupied areas and breakage regions devoid of cells. Fast Random Forest method was used considering gaussian blur, hessian, membrane projections, mean, entropy, sobel filter, difference in gaussians, Kuwahara and neighbors features and the following parameters: membrane thickness 1, membrane patch size 19, minimum sigma 1, maximum sigma 16. For the cell-cell junction, training was done by segmenting images into two classes: cell-cell junctions and the cytosolic area. Fast Random Forest method was used considering gaussian blur, hessian, membrane projections, mean, gabor, entropy, sobel filter, difference in gaussians, Kuwahara and neighbors features. Three sets of parameters have been used to account for the images’ brightness (faint, medium, bright). The parameters were: faint – membrane thickness 5, membrane patch size 25, minimum sigma 1, maximum sigma 20; medium – membrane thickness 5, membrane patch size 25, minimum sigma 1, maximum sigma 20; bright – membrane thickness 5, membrane patch size 25, minimum sigma 1, maximum sigma 20. This process generated three sets of binary images and a fourth image was obtained by combining the previous three binary images using the logic AND operator. These processes were performed via batch processing on ImageJ. Overlay of the segmented images on the original E-Cadherin image allowed us to retain the most accurate segmented image of this set, which was then converted into a 8-bit binary image. All binary images were then processed using morphological filters with MorpholibJ plugin ^16^ (closing operation, octagon element, and radius 4; directional filter with a max type, opening operation, line 26, and direction 32). Manual correction was performed on the images to eliminate any residual noise left during the segmentation process. Binary images were finally skeletonized to obtain the skeletonized image and were multiplied by the original E-Cadherin image to obtain a masked image where background pixels were removed while foreground pixels were retained. This process generated the following sets of images:

1 – Tissue breakage that was used to analyze the percentage of area devoid of cells (Figures 3 and 5).

2 – Cell-cell junction segmented, skeletonized, masked and nuclei image, that were used to build the classifier to distinguish tensional state of the tissues.

### Feature extraction

To build the tensional state classifier, a set of features was computed from the segmented, skeletonized, masked and nuclei images. All features were computed using Python and the libraries skan ^62^, OpenCV ^63^ and scikit-image ^64^

Several features were derived for each sample based on the four available images (Suppl. Table 2):

● Cell nuclei
● Original segmented cell-cell junction: cell-cell junctions are represented in white, while the remaining areas are in black.
● Masked cell-cell junction: the original pixels corresponding to the cell-cell junction are retained, with the remaining pixels rendered in black.
● Skeletonized cell-cell junction."

In the segmented images, features were calculated based on blobs, defined as clusters of black pixels. While these blobs often correspond to cells, this is not a requirement. Additionally, since an image may contain multiple blobs, it was necessary to aggregate the local descriptors to compute a global descriptor for the image. The median was used for this aggregation, unless differently specified. It is important to note that when aggregating the different blob descriptors, those located at the image edges were excluded.

Additionally, we computed features based on the image skeleton, which can be conceptualized as a graph consisting of nodes and edges, where each edge corresponds to a branch of the skeleton. There are three types of edges:

● Endpoint-to-endpoint (e2e): an isolated branch with no adjacent branches.
● Junction-to-endpoint (j2e): a terminal branch with an adjacent branch on only one side.
● Junction-to-junction (j2j): a branch with adjacent branches on both sides.

It is important to note that each branch is not necessarily a straight line; therefore, the length of a branch can be measured either by summing the pixels along the curve (referred to as "distance") or by calculating the Euclidean distance between the branch’s start and end points ("eu_distance").

### Machine learning and image classification

Experiments were designed to characterize the impact of nocodazole on tissue tension, leveraging the previously described and calculated features. The study employed control samples (NegsiRNA), including 34 samples treated with nocodazole (NO) and 34 samples without any stimulus (NS), to identify image features capable of differentiating between high tension (NO) and low tension (NS). With the features already computed, the initial step involved removing those that contributed minimal information, specifically eliminating features with zero variance or null values. Subsequently, 10 random forest models ^65^ were trained, each one with *n_estimators* inner decision trees (*n_estimators* ∈ [1500, …, 2500]), resulting in an average out-of-bag accuracy of 0.842 (± 0.004). The features were then ranked according to their importance as determined by the ensemble of random forests using Gini importance. Recognizing the potential for feature correlation, clusters were formed, with each cluster comprising features that exhibited an absolute Pearson correlation greater than T=0.8. From each cluster, only the feature with the highest importance was selected, reducing the set to 12 features.

In the subsequent step, a similar approach was employed, training independent 10 random forests with *n_estimators* again ranging from 1500 to 2500, but this time utilizing only the 12 uncorrelated features. This approach yielded an average out-of-bag accuracy of 0.854 (±0.004)). The features were again ranked by importance, and ultimately, only the two most significant features were retained. Those features were:

- *J2J length 1* (code name *j2j_distance_median*): a measure of the average length of the most common type of edge in the skeletonized image.

- *Junc. Thickness 1* (code name *branch_thickness_voronoi_median*): a measure of the width of each branch of the cell cell junction, computed using an approximation of the generalized Voronoi algorithm.

A linear Support Vector Machine (SVM) was trained using only these two features to classify samples as NS (low tension) or NO (high tension). The model was first evaluated using 10-fold cross-validation, achieving an accuracy of 0.826 (± 0.08), and then a final model was trained using all the data. This approach resulted in the creation of a 2-dimensional feature space, partitioned by a hyperplane according to the SVM’s classification. Subsequently, the remaining samples, which had knocked-down proteins, were projected into this vector space.

### Network of physical connections calculated from String

To build a network of the proteins positive to our screening, we employed STRING (https://string-db.org/). In this network we categorized proteins positive for our screening as interacting nodes in 4 distinct categories (Interface, Signaling, Cytoskeleton and Other functions, as in Suppl. Table 3) based on 3 criteria (as indicated in Figure 7): 1) Tissue disrupted > 5 % in homeostasis or upon induction of tension; 2) Machine learning classifier confidence > 90 % in homeostasis and/or upon induction of tension; 3) at least 2 positive hits with lower thresholds (i.e., confidence level between 60 and 90% and/or tissue breakage between 1 and 5% of the area) in homeostasis and/or upon induced tension. Only physical interactions (experiments) and no confidence threshold (score) were used (Suppl. Table 3).

### Cancer survival analysis using timer

Genes of proteins positive to our screening were evaluated through Kaplan–Meier survival analysis in TIMER ^20^ for the purposes of identifying specific genes whose changes in expression levels carry prognostic significance for cancer patient outcome (Figure 8 and Suppl. Table 5). z-scores comparing the survival rate of cancer patients with high versus low gene expression have been used to identify potential prognostic markers associated with mechanical effectors interfering tissue integrity.

### Statistics and Reproducibility

All representative figures reflect data extracted from three different biological replicates per protein KD, confidence intervals were calculated using quartiles (60%: Q.20 - Q.80, 90%: Q.05 - Q.95) obtained from Neg-siRNA control groups (n=34, per condition) using Google sheets, data visualization was made on Graphpad Prism 8.

## Acknowledgments

We are grateful to administrative staff at the Institute for Biological and Medical Engineering and at Faculty of Biology, the Unidad de Microscopia Avanzada (UMA) at Pontificia Universidad Catolica de Chile. We thank Daniela Salas and Daniel Navarro for efficient laboratory management.

## Fundings

Chilean Agency for Research and Development ANID SCIA ACT192015 (AR, CB, MC)

Chilean Agency for Research and Development ANID Fondequip EQM210101 (AR, CB)

Chilean Agency for Research and Development Fondecyt 1210872 (AR, CB)

Chilean Agency for Research and Development ANID Nucleo Milenio NCN2024_068 (AR, MC)

Chilean Agency for Research and Development ANID Fondequip EQM210020 (MC, CB)

Chilean Agency for Research and Development Fondecyt 1221696 (MC)

Chilean Agency for Research and Development Fondecyt 1230919 (MC)

Chilean Agency for Research and Development Fondecyt 1211988 (MC)

Pontificia Univ. Catolica de Chile Puente 2024-3 (CB)

National Research Foundation Singapore project NRF-MSG-2023-0001 (PK)

## Author contributions

Screening and image acquisition: CB and AS.

Image segmentation: SVS and RIP.

Machine learning classifier: JJA, MC, GM and AR.

Bioinformatic analysis: IAV, BCP, CSM, ATE and GIO.

Preparation of the figures: SVS, JJA and AR.

RNAseq: PK and CB.

Designed and supervised the study: RZB, CB and AR.

Wrote the manuscript: JJA, CB and AR.

All authors discussed the results and commented on the manuscript.

## Code Availability

The code for projecting samples into a two-dimensional space and computing the distance to the SVM-learned hyperplane is available in https://github.com/JuanjoAlegria/protein_analysis. The code for image feature extraction and feature selection is available upon request.

## Data Availability

All data supporting the conclusions are available from the corresponding author upon reasonable request.

---

**Suppl. Table 1.**
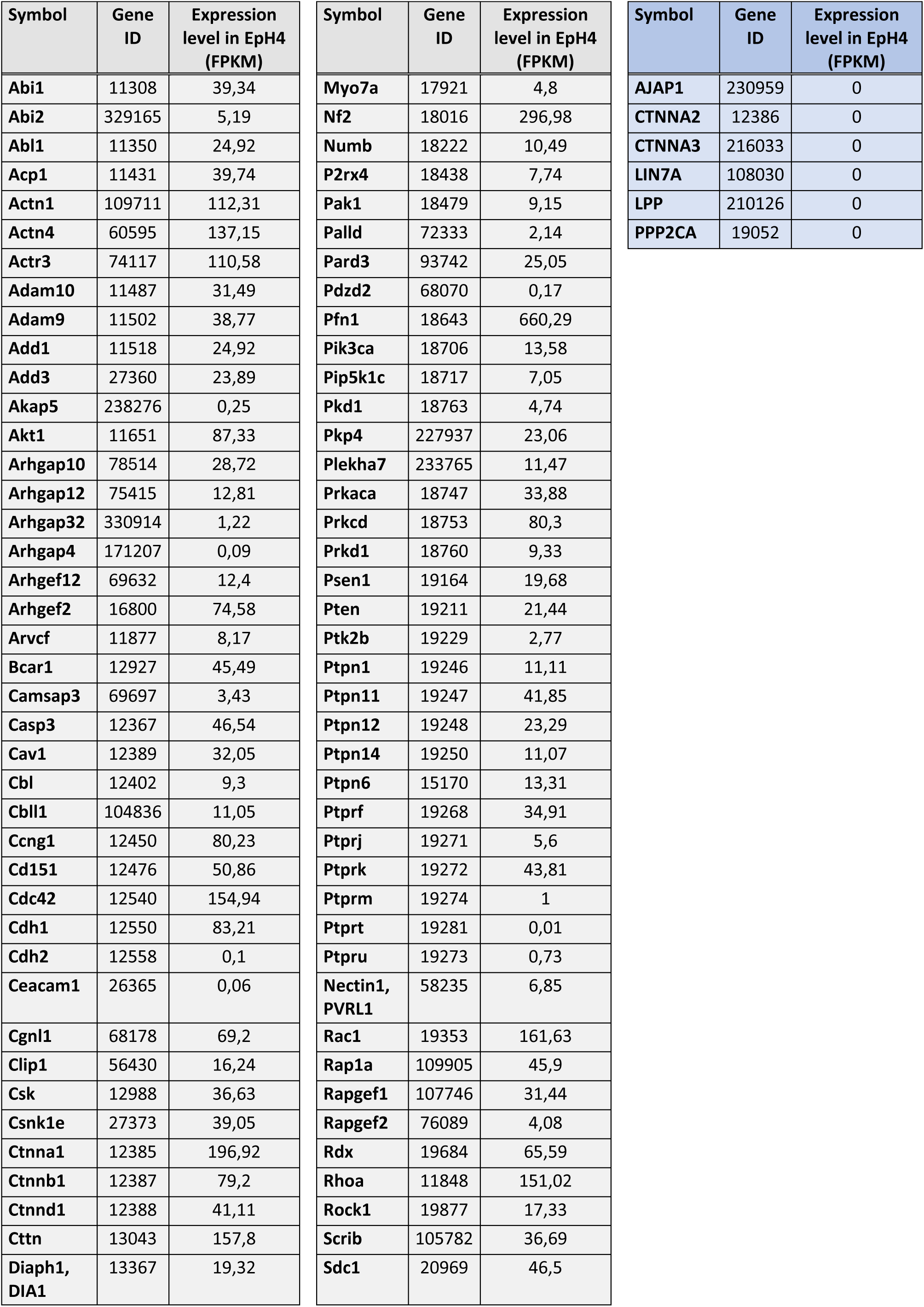
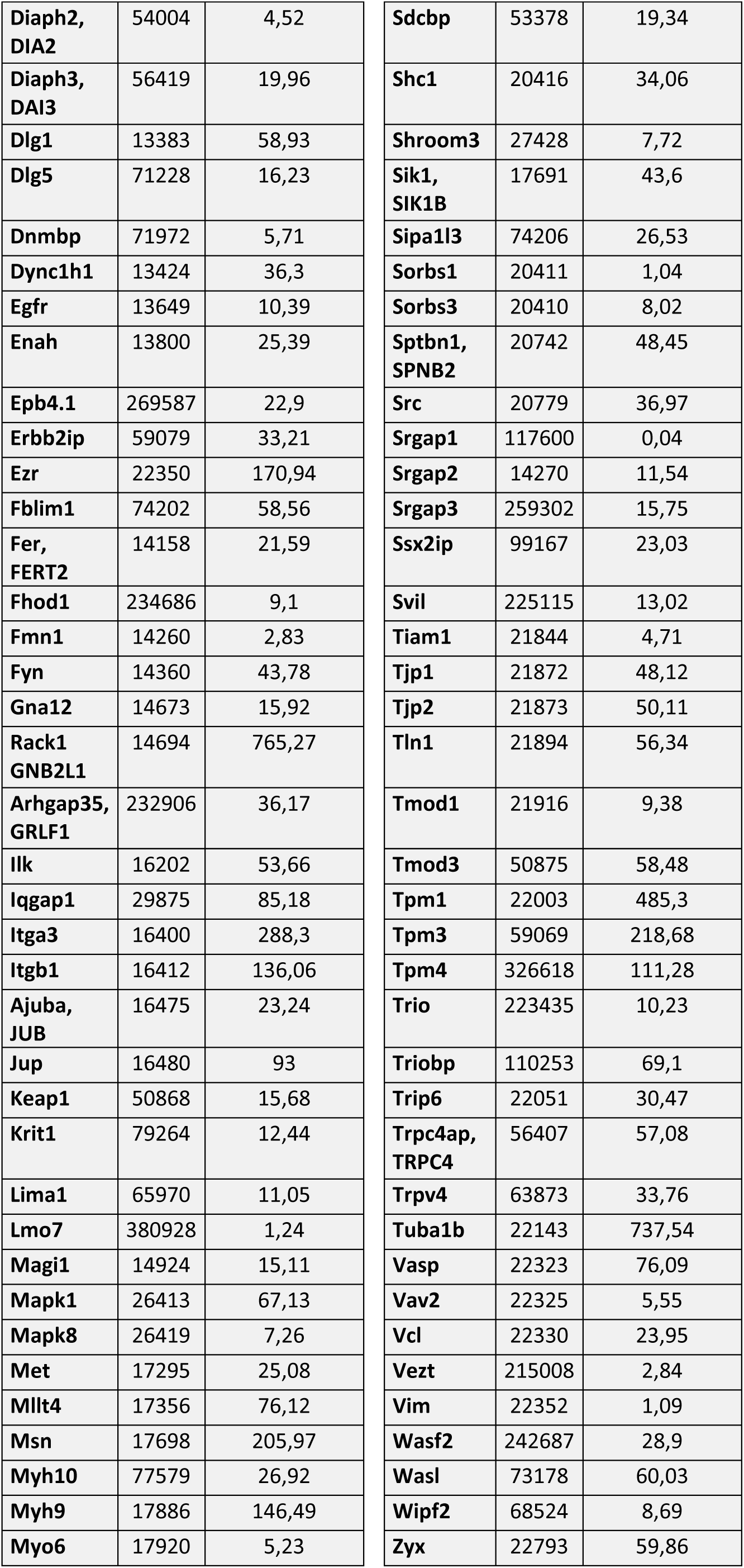
List of cadhesome genes expressed in EpH4 cells. Each column indicates the transcript ID, the Fragments Per Kilobase of transcript per Million mapped reads (FPKM), the gen Symbol in mouse, the description and the Blast number. The quantification of gene expression has been performed by mapping reads to the reference gene set that is Mus musculus (assembly GRCm38.p5).

**Suppl. Table 2.**
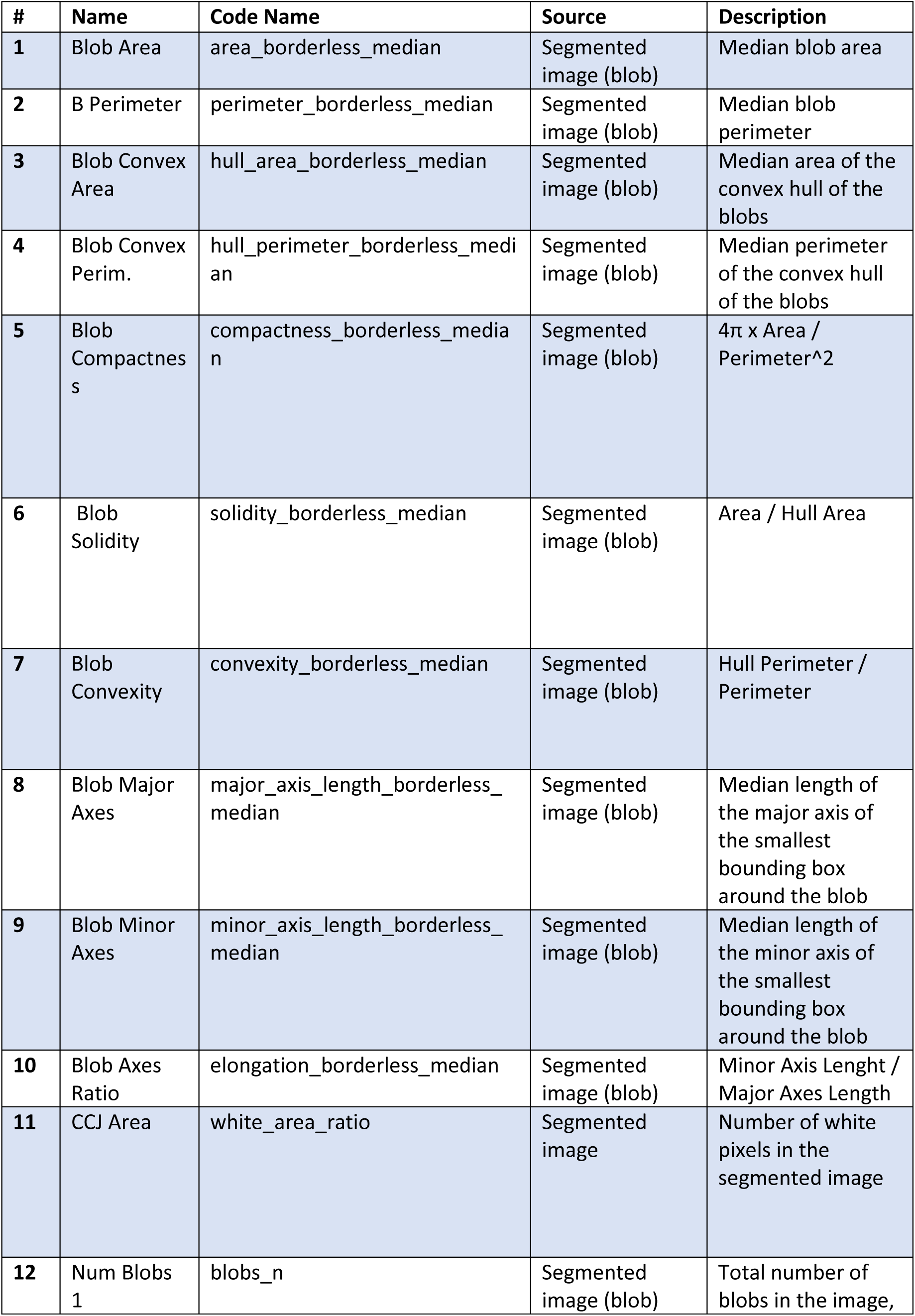
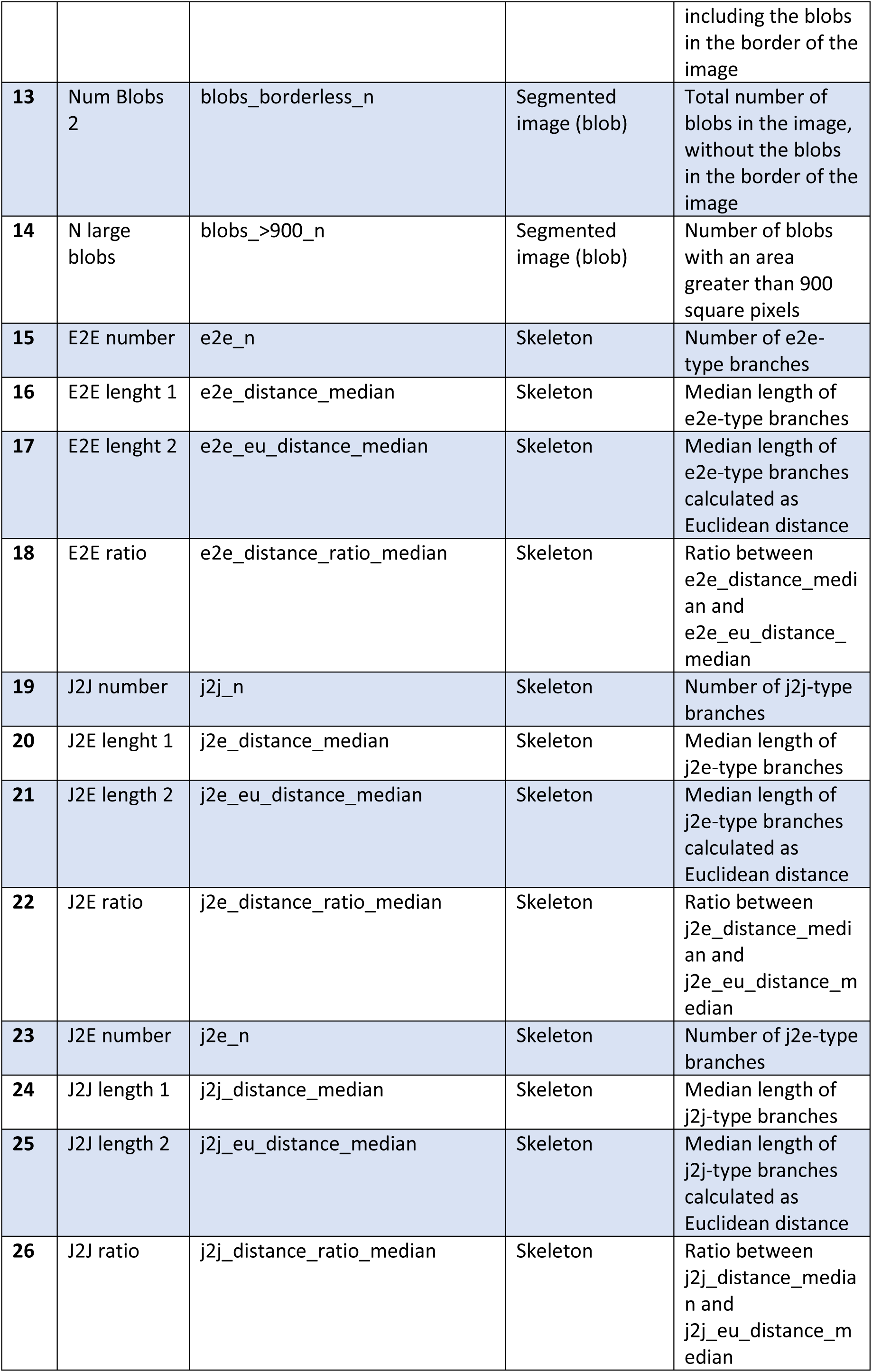
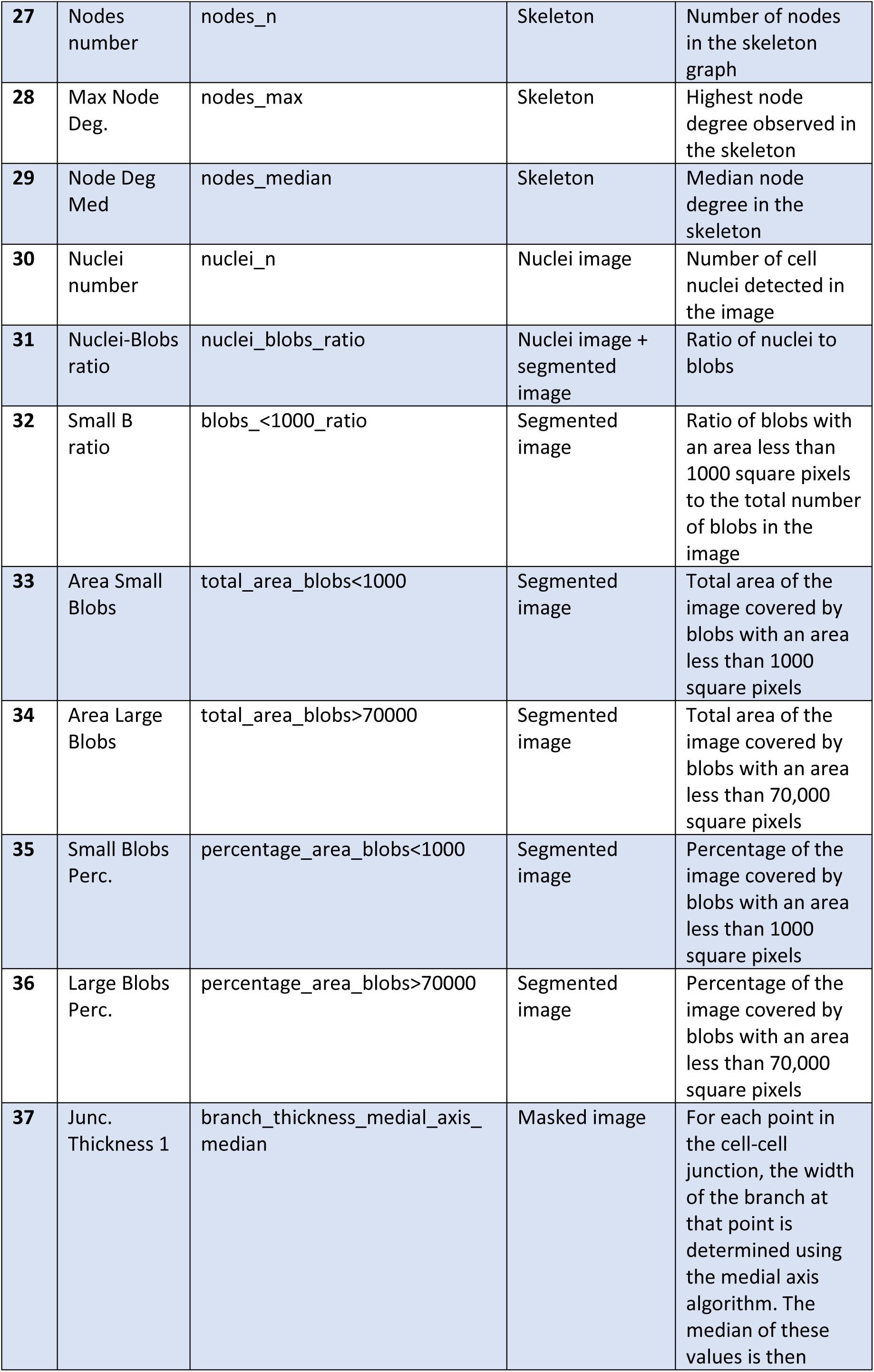
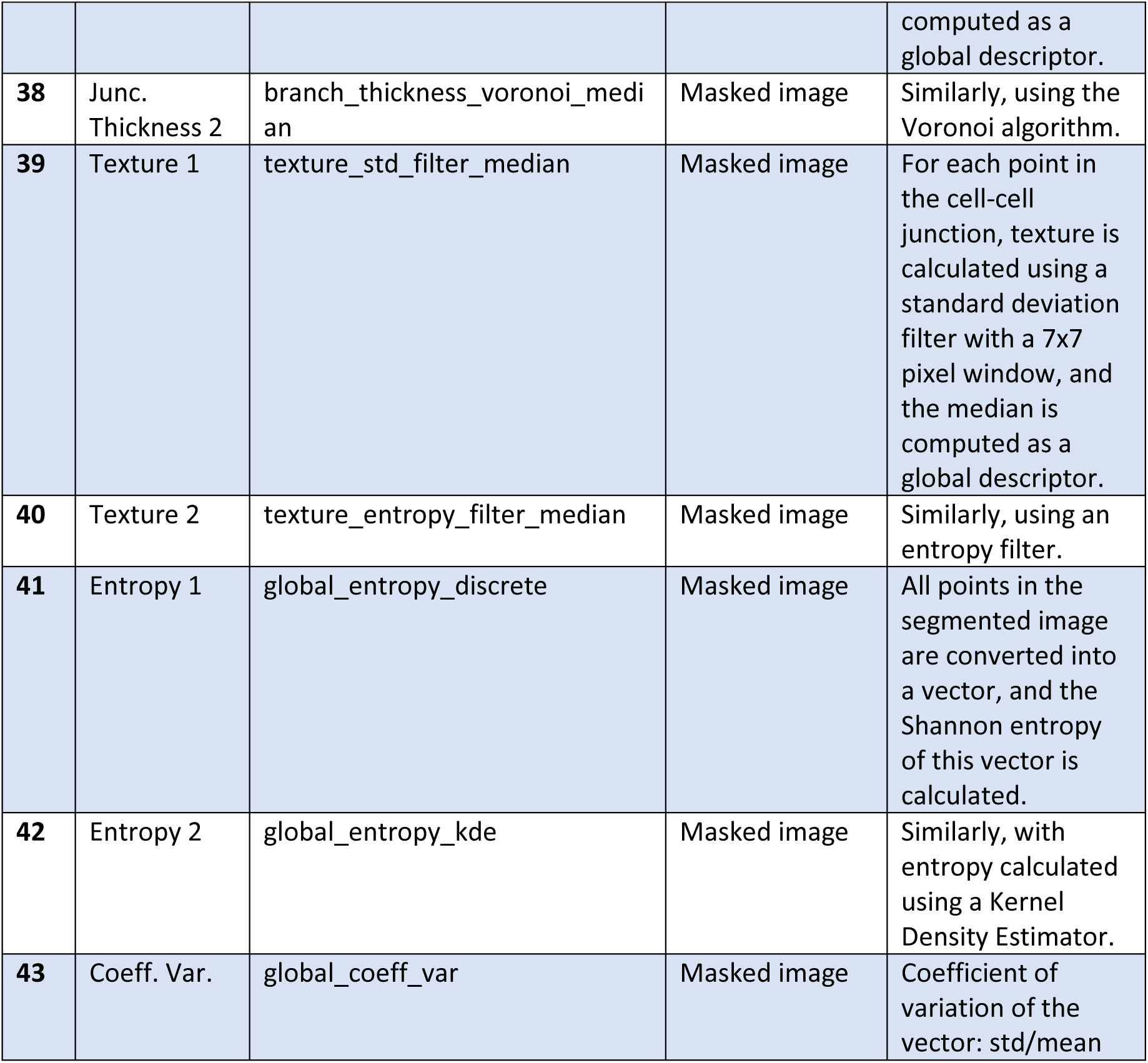
List of image features used in Machine Learning classifier. Different features extracted for each sample, specifying the type of image from which they were generated. Features highlighted in yellow passed the initial feature selection stage, meaning they had sufficient variance and no missing values (N=37, including blue features). Features highlighted in blue are those that were uncorrelated (N=12, including green features), and those in green represent the final features selected (N=2).

**Suppl. Table 3.**
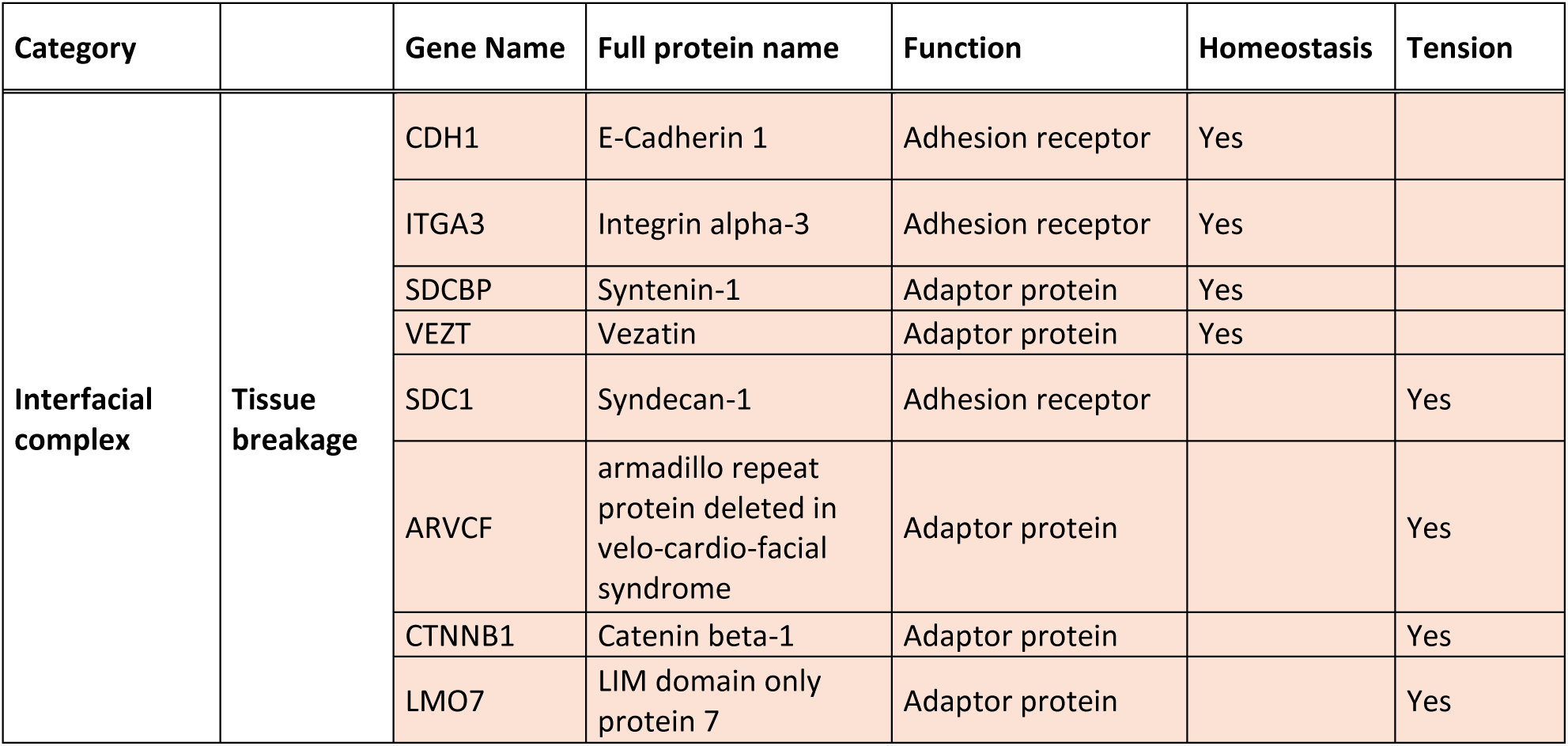
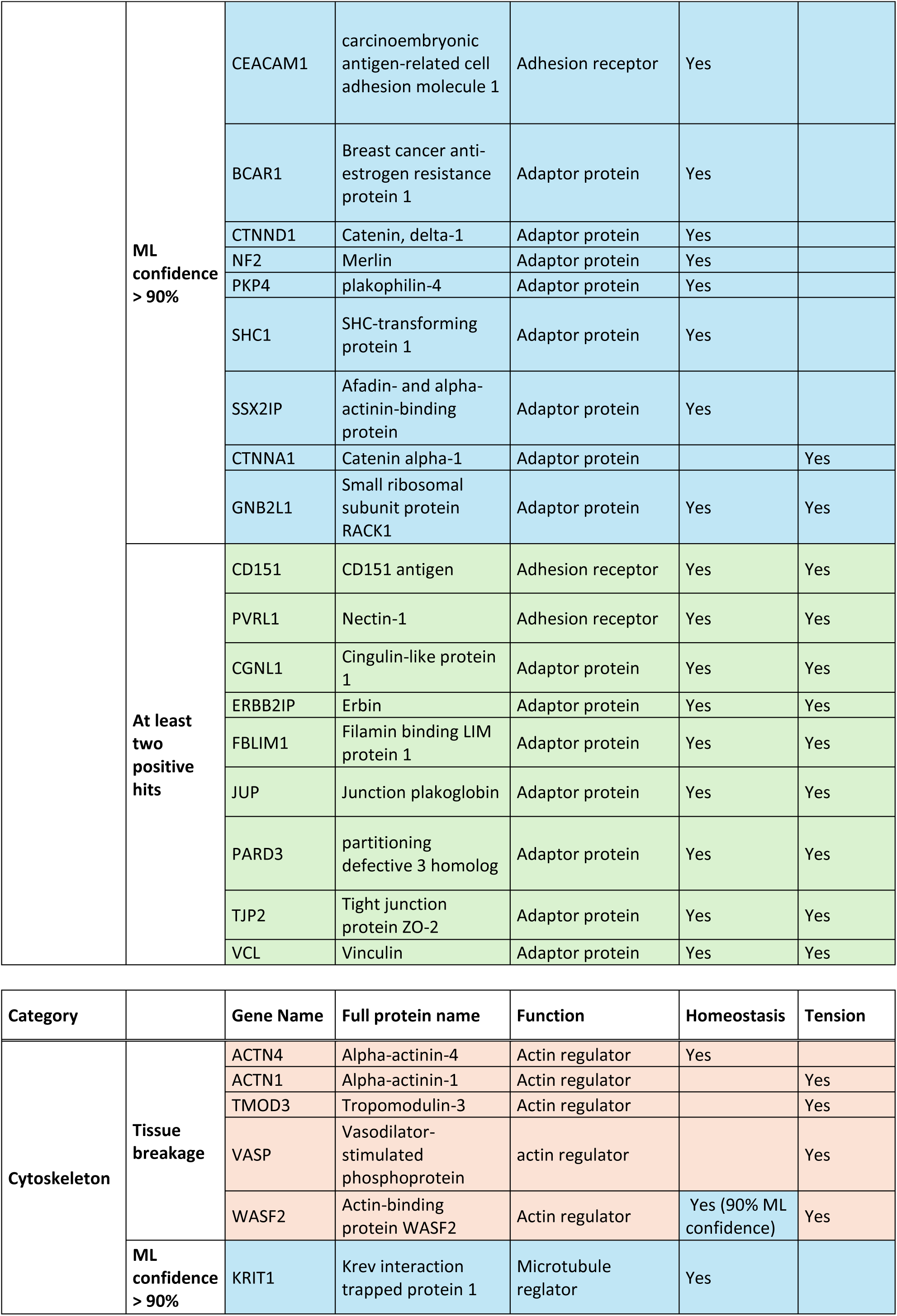
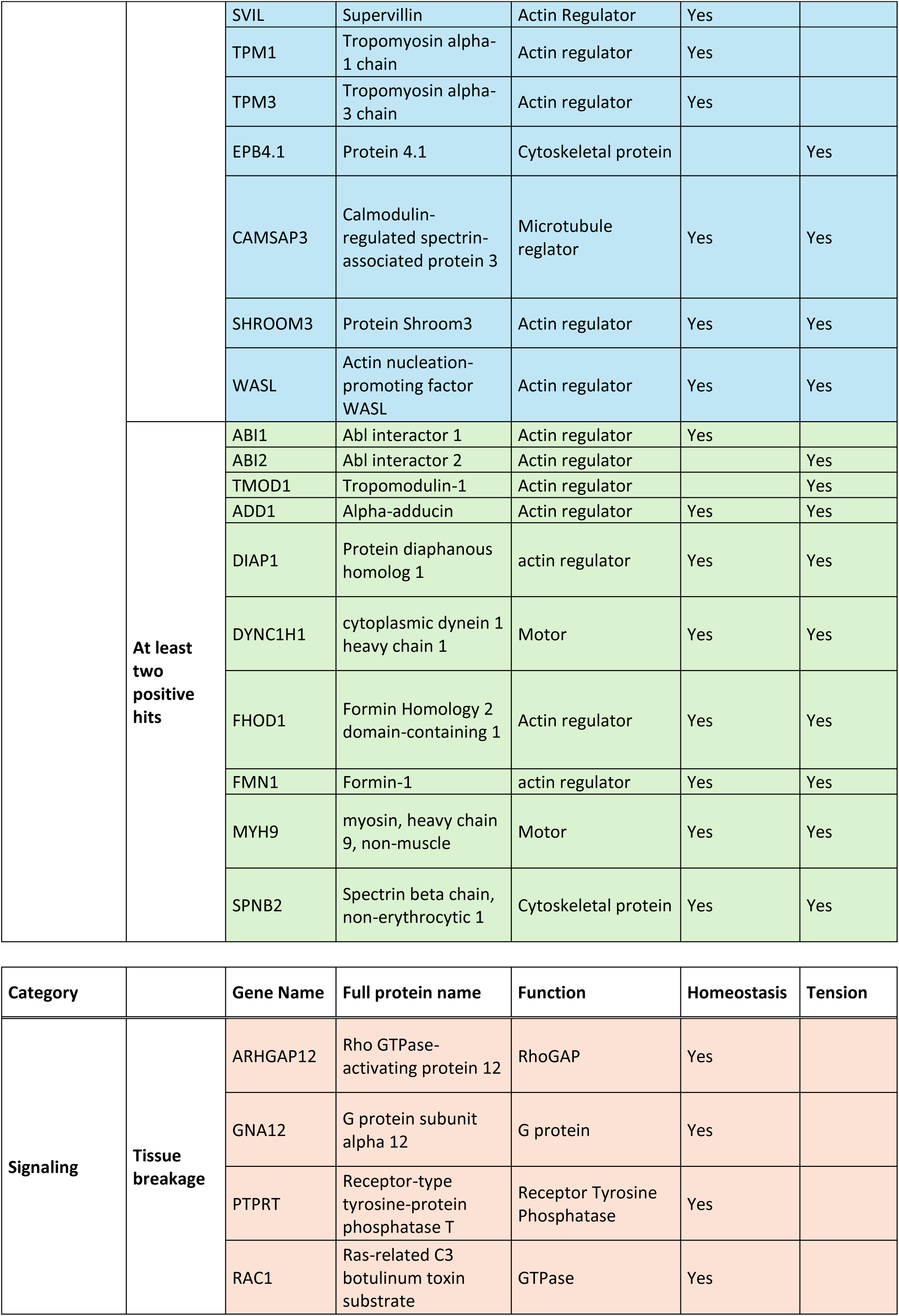
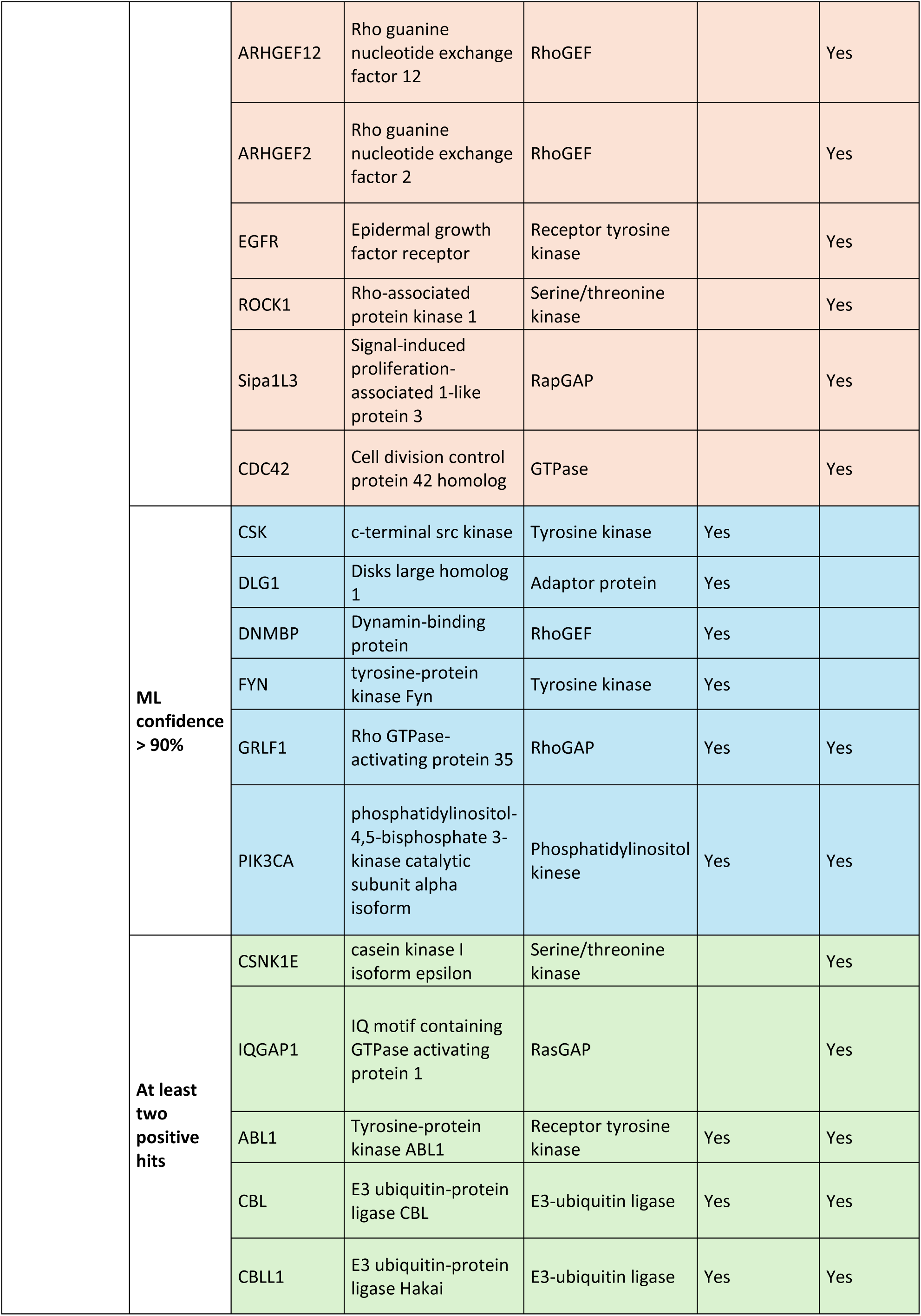
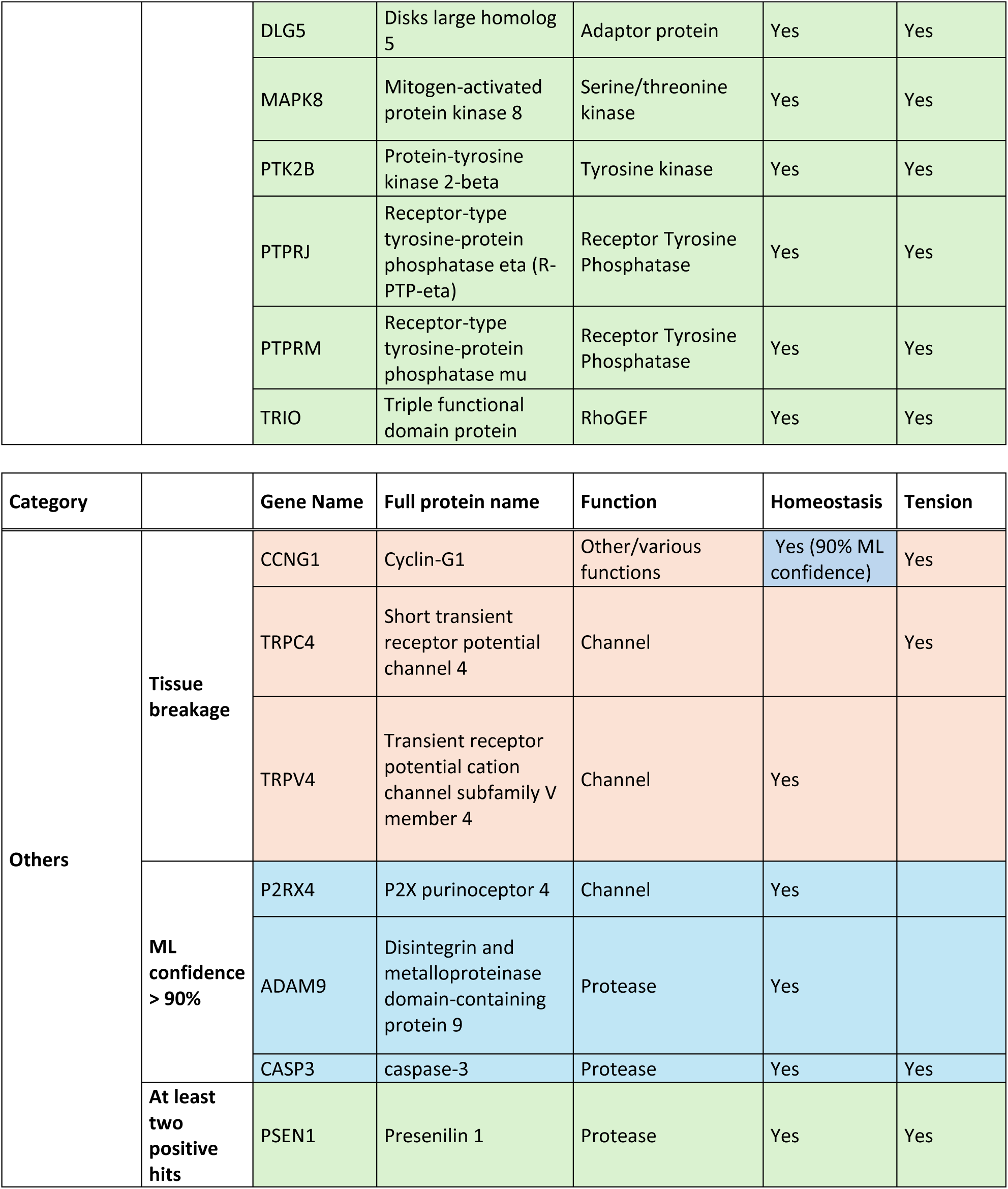
List of proteins positive of our screening subdivided into their respective functional categories.

**Suppl. Table 4.** STRING protein network (https://string-db.org/). Score reports STRING physical network derived from experiments.

| Node1 | Node2 | Score |
| --- | --- | --- |
| CTNNA1 | VCL | 0.965 |
| TPM1 | TPM3 | 0.963 |
| CDH1 | CTNNB1 | 0.962 |
| RAC1 | TRIO | 0.947 |
| CD151 | ITGA3 | 0.571 |
| CBL | CSK | 0.570 |
| ABI1 | ABL1 | 0.562 |
| ACTN4 | CTNNB1 | 0.550 |
| DIAPH1 | CDC42 | 0.388 |
| CASP3 | CDH1 | 0.386 |
| PARD3 | CTNNA1 | 0.372 |
| ADD1 | TPM3 | 0.371 |
| CDH1 | JUP | 0.943 |
| ACTN1 | ACTN4 | 0.933 |
| ACTN1 | VCL | 0.902 |
| VCL | ACTN4 | 0.902 |
| ABI2 | RAC1 | 0.869 |
| CDH1 | GNB2L1 | 0.833 |
| CBL | EGFR | 0.788 |
| CTNNA1 | CTNNB1 | 0.787 |
| CTNNA1 | JUP | 0.770 |
| ABI1 | WASF2 | 0.738 |
| EGFR | SHC1 | 0.681 |
| EGFR | CSK | 0.677 |
| CTNND1 | CTNNB1 | 0.668 |
| JUP | CTNNB1 | 0.657 |
| ABI2 | WASF2 | 0.651 |
| CTNND1 | CDH1 | 0.647 |
| ABI1 | ABI2 | 0.636 |
| CDC42 | IQGAP1 | 0.631 |
| CBL | SHC1 | 0.625 |
| DNMBP | WASL | 0.618 |
| BCAR1 | VCL | 0.617 |
| CSK | SHC1 | 0.615 |
| ARVCF | CTNNB1 | 0.614 |
| WASL | WASF2 | 0.614 |
| RAC1 | IQGAP1 | 0.604 |
| PARD3 | CTNNB1 | 0.550 |
| PIK3CA | EGFR | 0.550 |
| ACTN1 | CTNNA1 | 0.548 |
| CDC42 | MAPK8 | 0.542 |
| FYN | CTNNB1 | 0.542 |
| FYN | CBL | 0.537 |
| FYN | CSK | 0.535 |
| PKP4 | CTNNB1 | 0.530 |
| ABL1 | ABI2 | 0.529 |
| ABL1 | EGFR | 0.523 |
| CTNND1 | EGFR | 0.523 |
| MAPK8 | EGFR | 0.523 |
| WASL | EGFR | 0.523 |
| ARHGEF12 | GNA12 | 0.515 |
| BCAR1 | FYN | 0.508 |
| ARHGAP12 | WASL | 0.495 |
| GFRL1 | RAC1 | 0.483 |
| CBLL1 | CDH1 | 0.483 |
| PTPRJ | CTNND1 | 0.483 |
| PIK3CA | CBL | 0.482 |
| NF2 | EGFR | 0.476 |
| CDC42 | WASL | 0.458 |
| GFRL1 | CDC42 | 0.457 |
| BCAR1 | PTK2B | 0.457 |
| ERBB2IP | PKP4 | 0.457 |
| FHOD1 | RAC1 | 0.457 |
| IQGAP1 | CTNNB1 | 0.457 |
| MAPK8 | CBL | 0.457 |
| PSEN1 | CDH1 | 0.457 |
| PSEN1 | CTNNB1 | 0.457 |
| PSEN1 | CTNND1 | 0.457 |
| PTK2B | CBL | 0.457 |
| PTK2B | EGFR | 0.457 |
| PTPRM | GNB2L1 | 0.457 |
| PTPRM | CDH1 | 0.457 |
| GNB2L1 | FYN | 0.457 |
| GNB2L1 | PTPRM | 0.457 |
| VASP | VCL | 0.457 |
| DNMBP | CDC42 | 0.456 |
| TPM1 | ACTN1 | 0.450 |
| ERBB2IP | CTNNA1 | 0.432 |
| ADD1 | TMOD1 | 0.422 |
| ARHGAP12 | WASF2 | 0.422 |
| ARHGAP12 | ABI1 | 0.420 |
| CASP3 | ROCK1 | 0.420 |
| DLG5 | CTNNB1 | 0.420 |
| PTPRJ | CTNNB1 | 0.369 |
| ARVCF | ERBB2IP | 0.365 |
| VASP | CDC42 | 0.363 |
| SVIL | ADD1 | 0.359 |
| ARHGEF2 | RAC1 | 0.316 |
| ADAM9 | CDH1 | 0.292 |
| ARHGEF12 | EGFR | 0.292 |
| CBLL1 | CDC42 | 0.292 |
| CEACAM1 | SHC1 | 0.292 |
| CSNK1E | CTNNB1 | 0.292 |
| DLG1 | PIK3CA | 0.292 |
| DYNC1H1 | EPB41 | 0.292 |
| EPB41 | TJP2 | 0.292 |
| GNA12 | CDH1 | 0.292 |
| MYH9 | ROCK1 | 0.292 |
| PVRL1 | CBL | 0.292 |
| NF2 | SDCBP | 0.292 |
| PTPRM | CTNND1 | 0.292 |
| SDC1 | SDCBP | 0.292 |
| SSX2IP | ACTN1 | 0.292 |
| TJP2 | CTNNA1 | 0.292 |
| TRPV4 | FYN | 0.292 |
| VEZT | CDH1 | 0.292 |
| TMOD3 | SVIL | 0.230 |
| FHOD1 | CDC42 | 0.229 |
| LMO7 | ACTN4 | 0.214 |
| TMOD1 | TPM1 | 0.211 |
| FBLIM1 | VASP | 0.187 |
| SHROOM3 | VASP | 0.165 |
| SIPA1L3 | CTNNB1 | 0.165 |
| TRIO | CDC42 | 0.165 |
| CGNL1 | CTNNB1 | 0.163 |
| ITGA3 | ABI1 | 0.161 |
| CAMSAP3 | CGNL1 | 0.041 |
| CD151 | CDH1 | 0.041 |
| FMN1 | CTNNA1 | 0.041 |
| KRIT1 | CTNNB1 | 0.041 |
| TRPC4 | TRPV4 | 0.041 |
| TPM3 | ACTN1 | 0.420 |
| PTPRJ | EGFR | 0.412 |
| PVRL1 | SDCBP | 0.404 |
| DLG5 | TPM1 | 0.402 |
| MYH9 | CBL | 0.400 |

**Suppl. Table 5.**
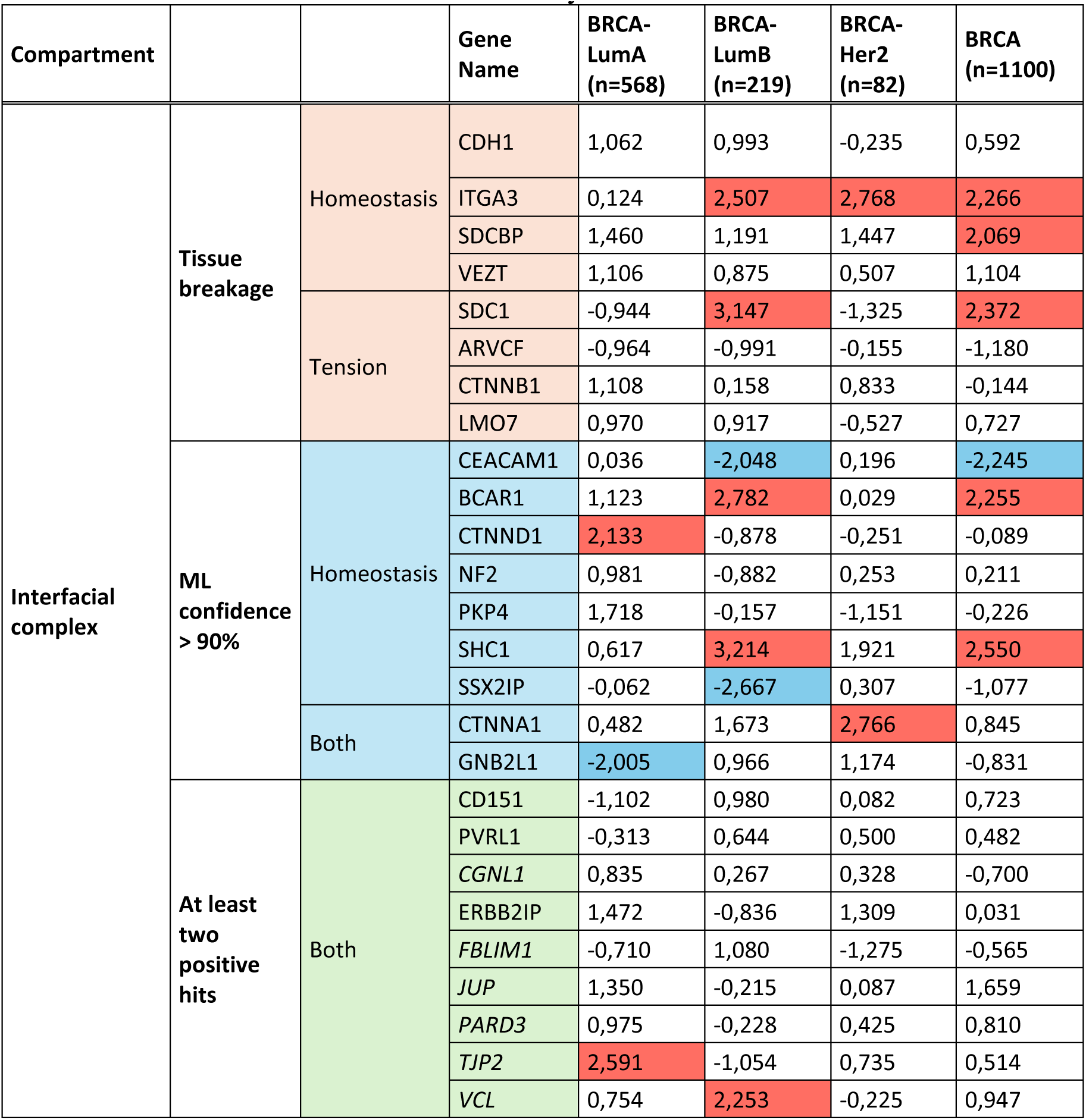
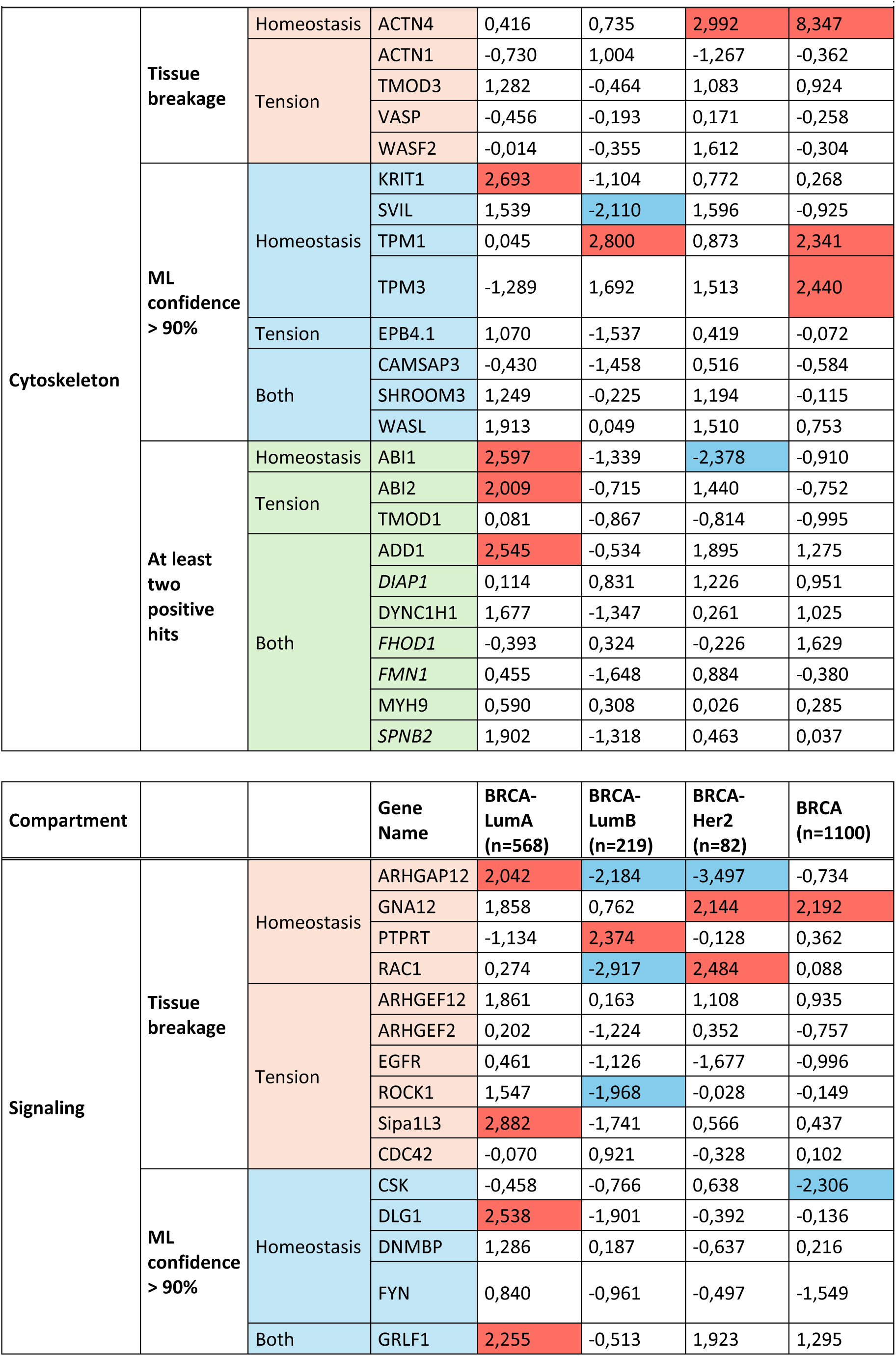
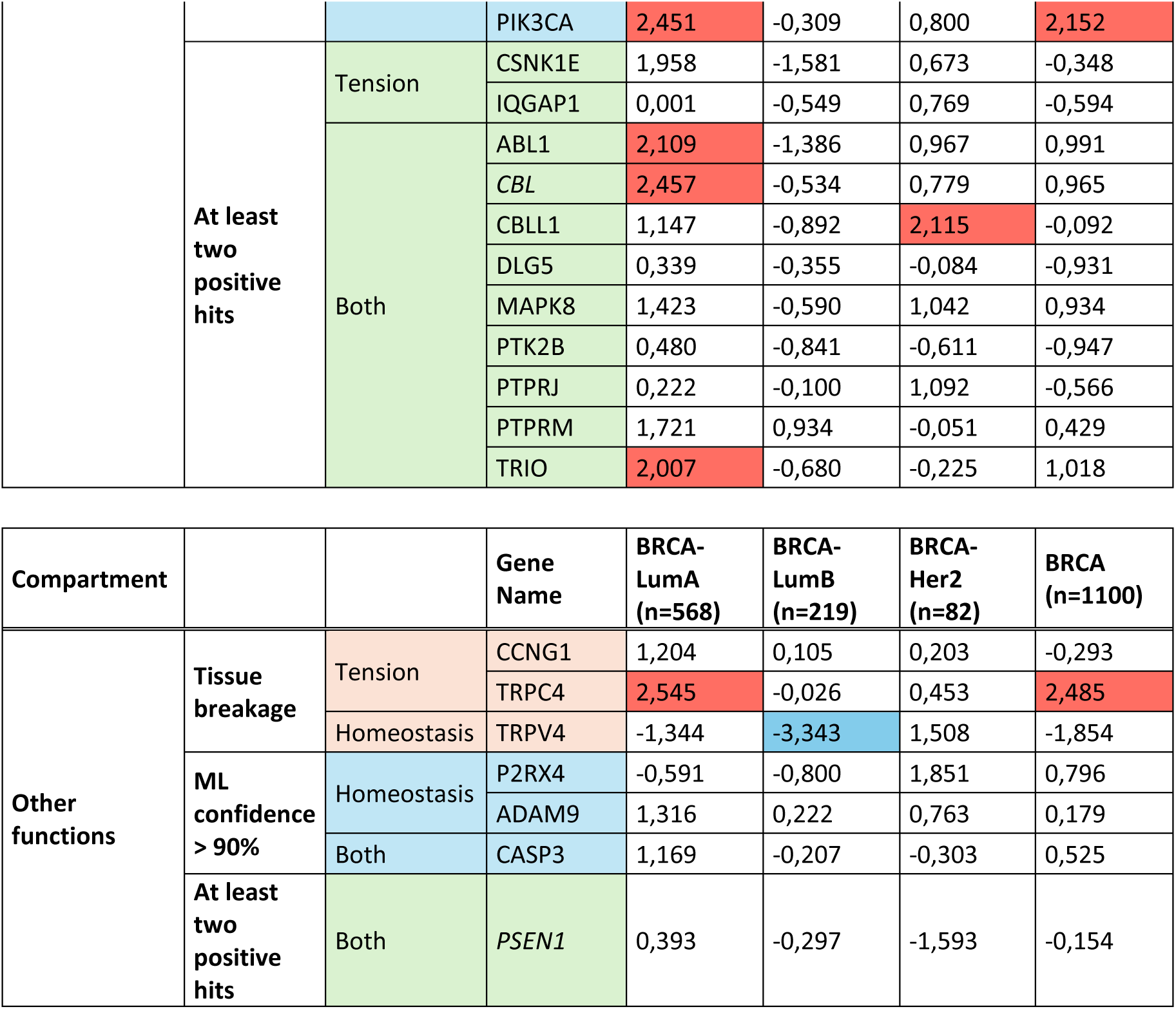
Gene hits analysis based on Kaplan–Meier survival of cancer patient obtained from TIMER. Table present z-scores comparing breast cancer patient survival rates as analyzed using the Kaplan– Meier Data are taken from the online resource TIMER. Positive z-score indicates that high patient survival is associated with low gene expression, which is consistent with our gene silencing screening, whereas a negative z-score (blue) indicates that low survival correlates with low gene expression. Statistically significant results (p < 0.05) are highlighted in red and blue backgrounds for high and low survival, respectively. In this test, the null hypothesis assumes that gene expression levels do not impact patient survival, and small p-values support rejecting this hypothesis. Cancer types and sample sizes are indicated in brackets under their acronyms in the table.

**Suppl. figure 1.**
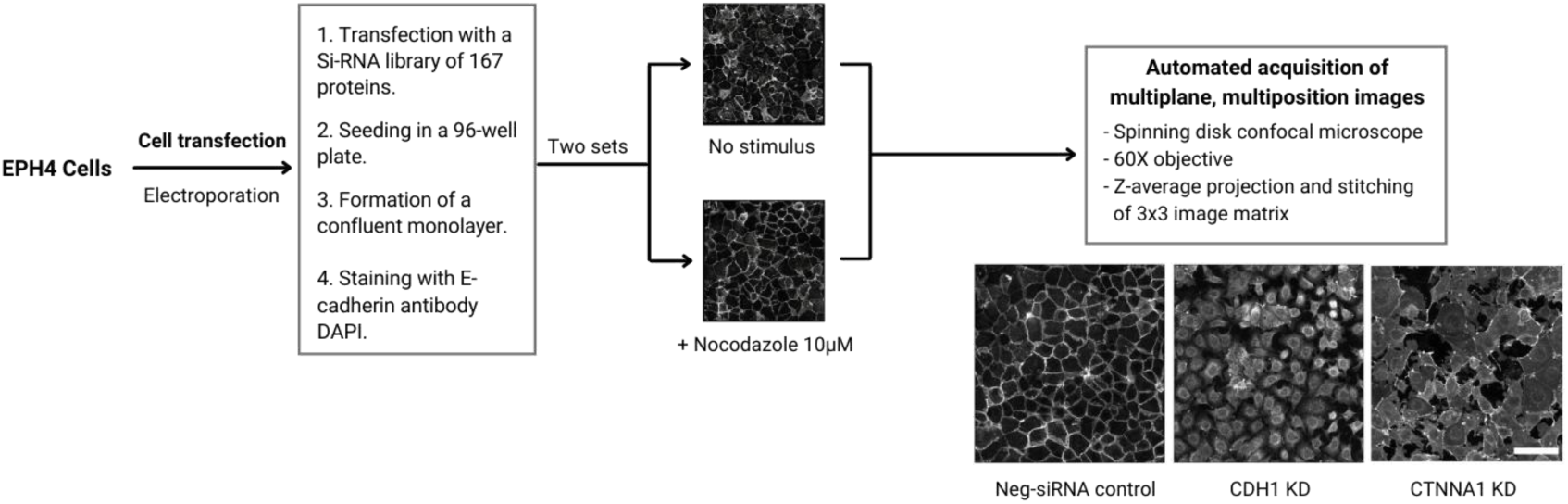
Generation of the knockdown library for proteins of the cadhesome and screening. Batches of suspended EPH4 cells were transfected with one GeneX-siRNA construct from a library of 167 proteins or with a Neg-siRNA scrambled sequence as a control for wild-type tissue. Cells were then seeded in repeats of 6 and incubated for 24 hours to form a monolayer. Three of these repeats were treated with nocodazole to induce tension within the tissue. The imaging process of the resulting cell monolayers was performed using a NIKON spinning disk confocal microscope and implementing a semi-automated acquisition protocol in a 3x3 grid. Scale bar = 50 µm.

**Suppl. Figure 2.**
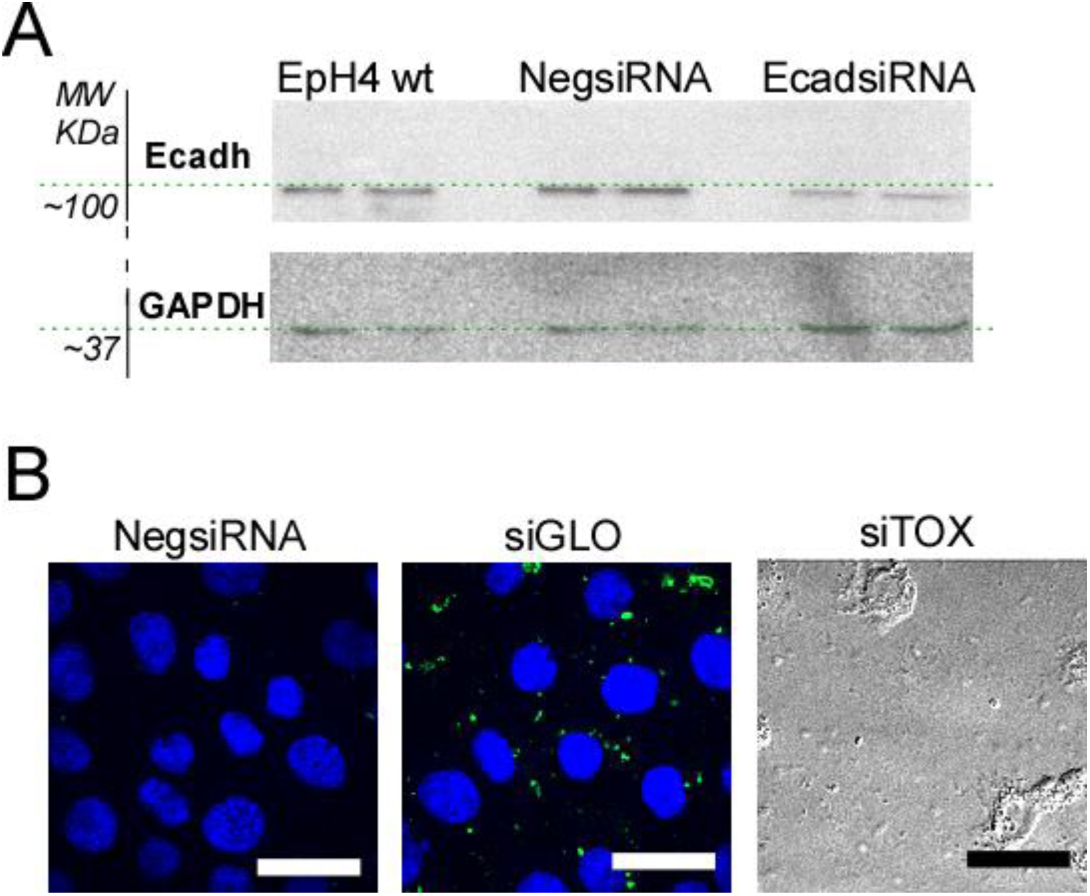
Validation of Transfection. **A**) Validation of si-RNA silencing of specific proteins was performed by Western Blot. Exemplary WB of E-Cadherin expression level for cells transfected by siRNA SMARTpool targeting CDH1, to validate our transfection efficiency. Expression level of E-Cadherin and GAPDH as loading control, in EpH4 cells wt, transfected with non-target siRNA (Neg-SiRNA), and transfected by the respective siRNA SMARTpool targeting CDH1. **B**) Each 96-well plate contained specific controls: samples transfected with non-target siRNA (Neg-SiRNA), which served as control for wild type tissue and two samples transfected with the transfection indicators, siTOX Transfection Control, and with siGLO RISC-Free Control siRNA, respectively, as control for transfection efficiency. Cell successfully transfected with siGLO shown appearance of siRNA spots (in green in the middle), while transfection with siTOX induced apoptosis and cell death within 24hours. Images were acquired by an inverted microscope (Eclipse Ti, Nikon, Japan) using a Nikon a Plain Apo Vc 60X oil objective, N.A. 1.40. Scale bar = 20µm.

**Suppl. Figure 3.**
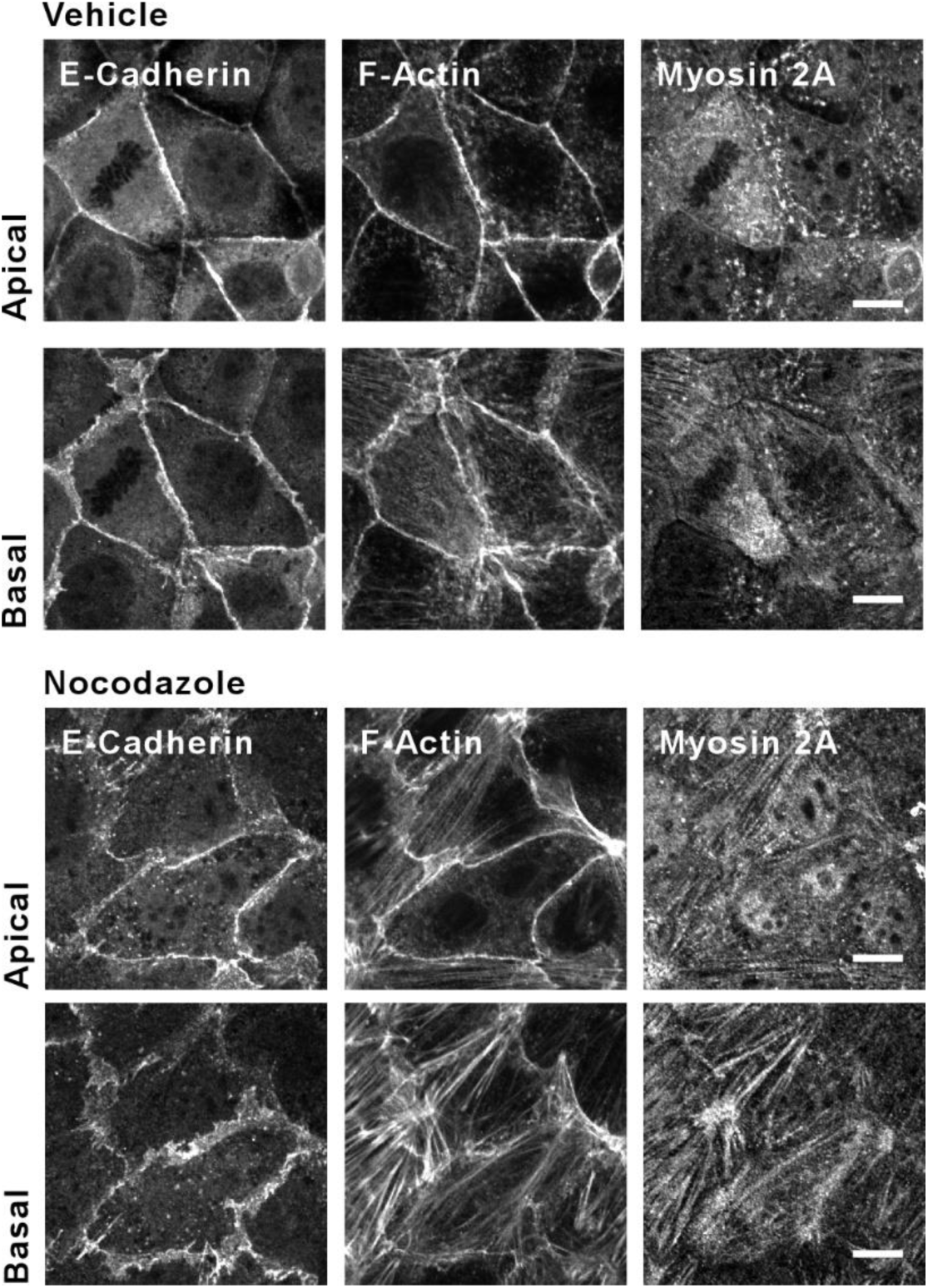
Validation of induction of tension by chemical treatment. EpH4 cells non treated (vehicle) and chemically treated (Nocodazole), stained for Ecadherin, F-actin and Myosin IIA. Imaging at different focal plane to highlight acto-myosin stress fibers (Basal), junctional actin and myosin organization (Apical). Images have been acquired by a CSU spinning disk (Yokogawa®) confocal head mounted on an inverted microscope (Eclipse Ti, Nikon, Japan) using a Plain Apo Vc 60X oil objective, N.A. 1.40. Scale bar = 5um.

**Suppl. Figure 4.**
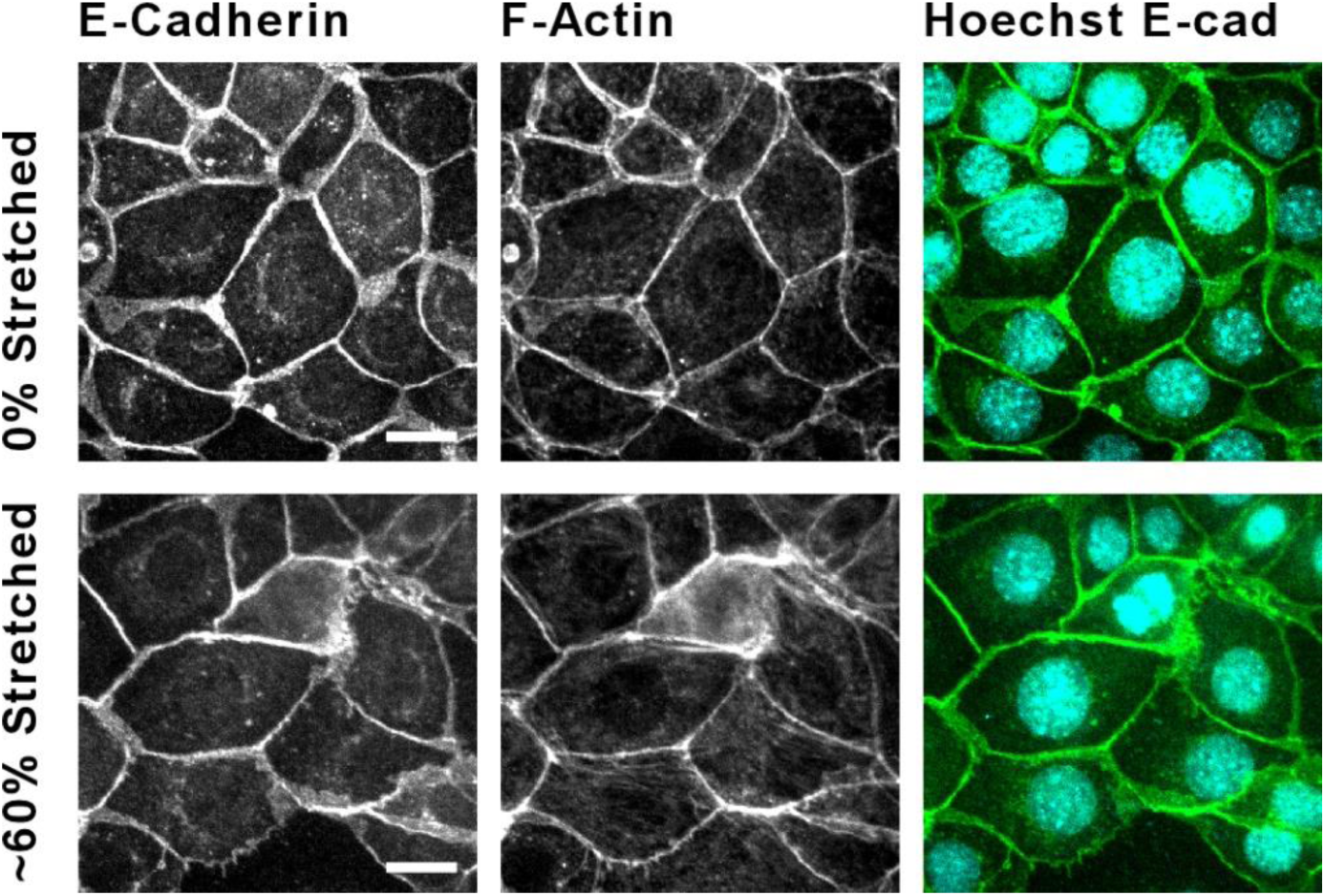
Mechanical induction of Tension. EpH4 cells non stretched (0% stretched) and mechanically stretched (60% stretched), stained for Ecadherin, F-actin and Hoechst. Images clearly highlight increase in stress fibers appearance, thickening and extension of cadherin-based cell-cell adhesions with increasing stretch. Images have been acquired by a CSU spinning disk (Yokogawa®) confocal head mounted on an inverted microscope (Eclipse Ti, Nikon, Japan) using a Plain Apo Vc 60X oil objective, N.A. 1.40. Scale bar = 5µm.

**Suppl. Figure 5.**
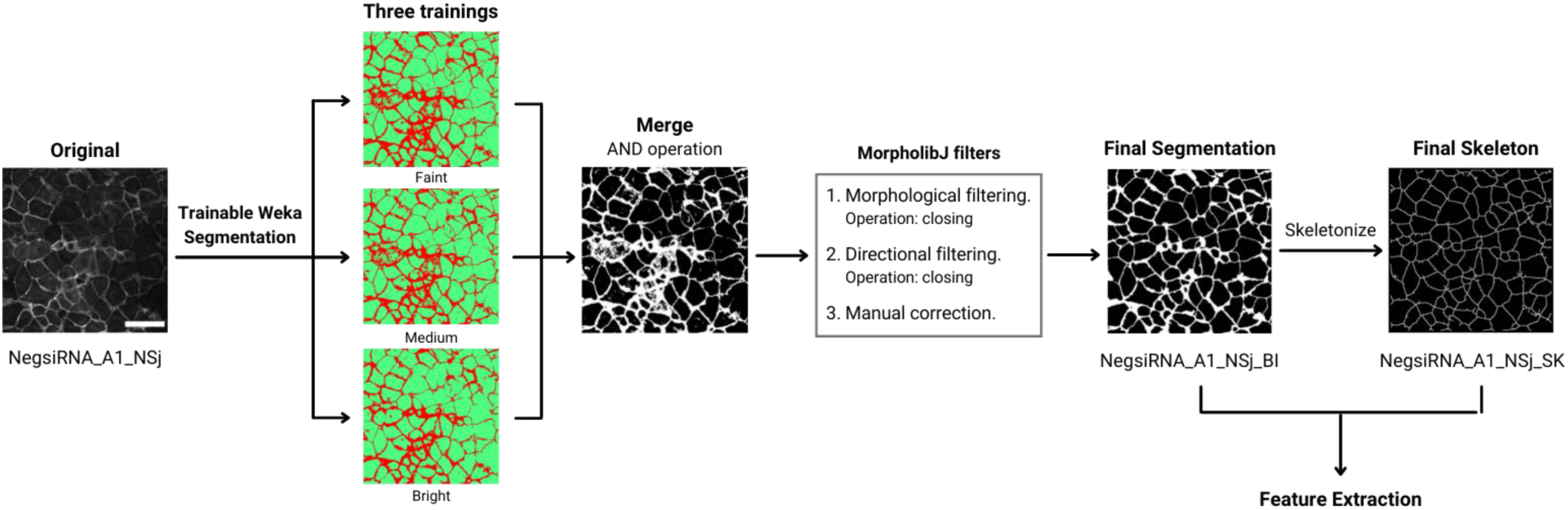
Image segmentation was based on Fast Random Forest using the Weka segmentation plugin of ImageJ. Three independent trainings were performed by selecting 20 representative images from the dataset, resulting in three separate segmentations, bright, medium, and weak, which were then converted to binary and combined using the AND logic operator. After merging, sequential morphological and directional filters were applied, consisting of a closing, opening, and closing operation from the MorpholibJ plugin of ImageJ to reduce noise and smaller holes. The final step consisted of manual correction of the images to guarantee the quality of the segmentation. These segmented images, in combination with a skeletonized version of them, were used for feature extraction in later steps. Scale bar = 50 µm.

**Suppl. Figure 6.**
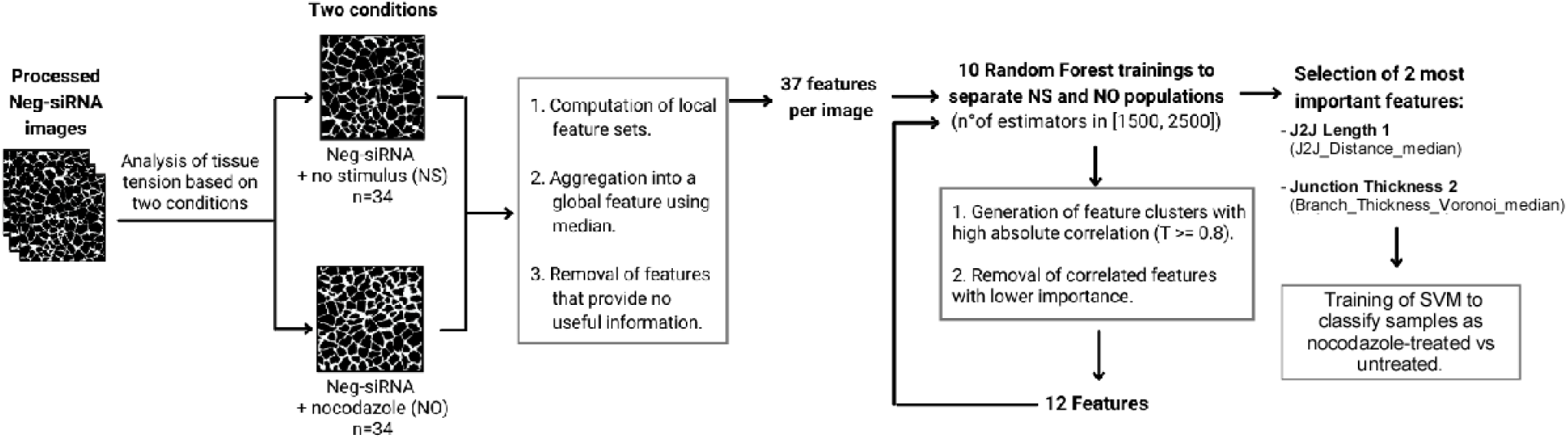
Classification pipeline to classify samples as nocodazole-treated vs untreated. The dataset comprised 68 images (34 nocodazole-treated and 34 untreated). Multiple features were computed for each image; since some were local features (i.e., representing only a portion of the image), these were aggregated by the median to derive a global feature representing the entire image. Features with minimal variation across images were excluded, resulting in a set of 37 features (Suppl. Table 2). Using these 37 features, 10 random forest models were trained to classify samples as nocodazole-treated versus untreated. After model training, the 37 features were ranked by their importance in the random forest models. Clusters of features with high absolute correlation (T > 0.8) were identified, retaining only the most important feature in each cluster and discarding the rest, which reduced the set to 12 features. This process was repeated once more, yielding two final features: junction length (code name j2j_distance_median) and junction thickness (code name branch_thickness_voronoi_median). A final linear SVM model was trained using these two features to distinguish between nocodazole-treated and untreated samples, achieving an accuracy of 0.826. This trained SVM model can subsequently be used to differentiate images by low(er) versus high(er) tensional state.

**Suppl. Figure 7.**
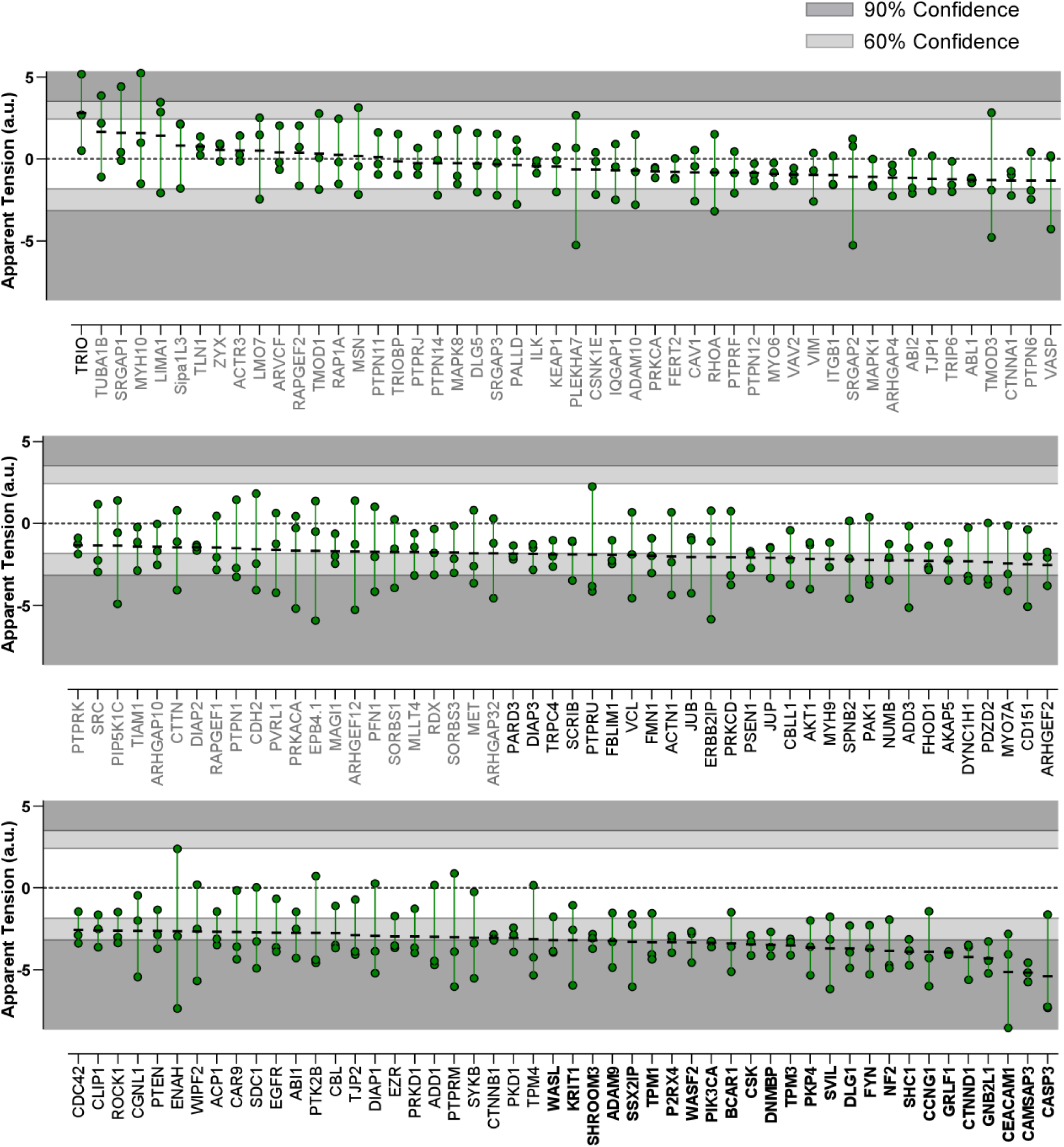
Identification of Proteins Playing a Role in Regulating Tensional State of the Tissue. Replot of data as in Figure 4 displayed as grouped data scatter plot of individual values. Individual measurements (repeats from different experiments, n=3 for each GeneX-siRNA) are depicted by green circles with measurement range indicated by the connecting green line and the mean of three independent measurements by a black dash. Dark and light grey areas indicate the confidence level of 60 and 90%, respectively. Protein names are in black bold font for confidence level above 90%, in black font for levels between 60 and 90 % and gray fonts for levels below 60%. Only samples whose % of breakage is below 5% are analyzed using this classification method (Figure 3).

**Suppl. Figure 8.**
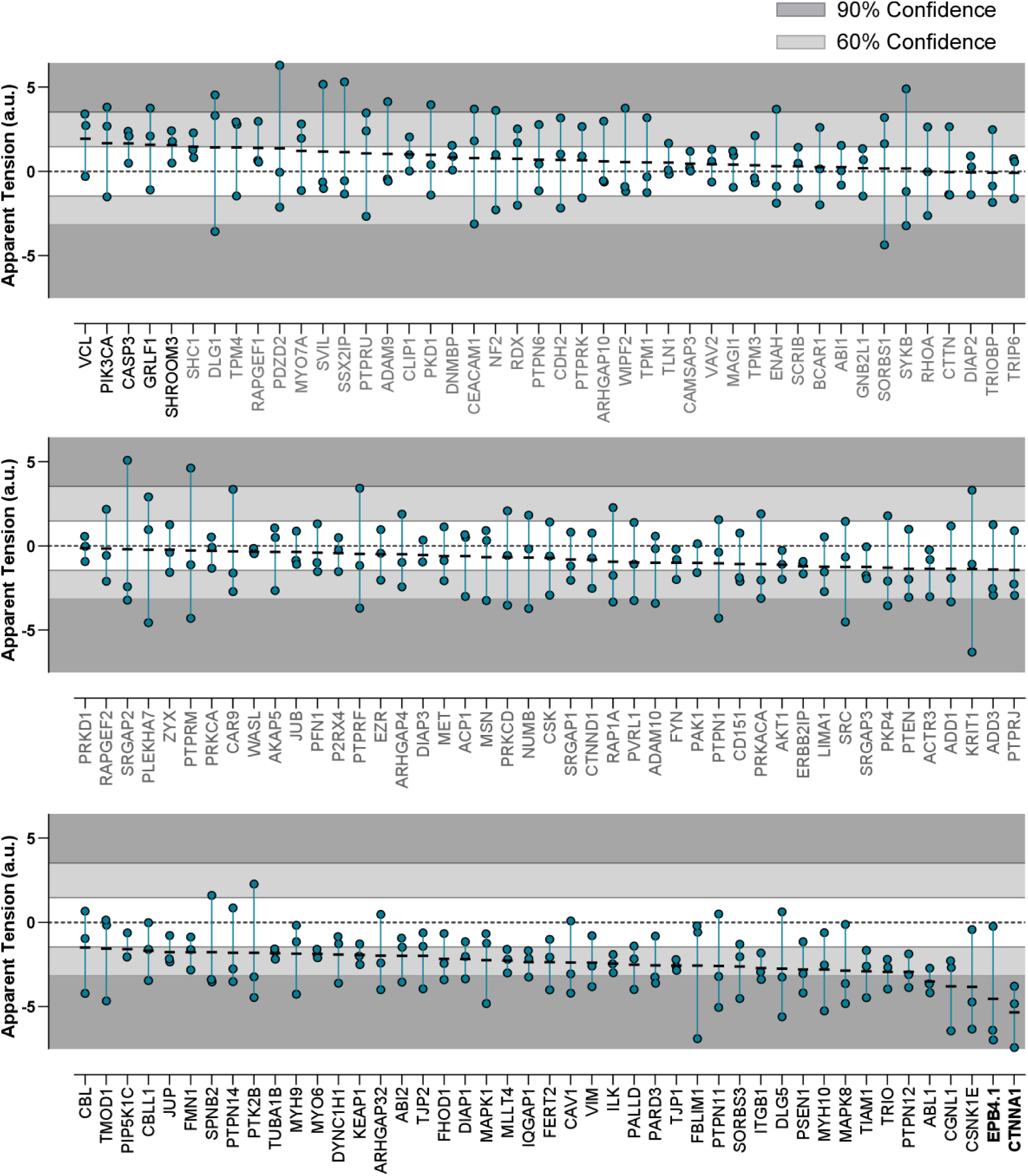
Identification of Proteins Playing a Role in Regulating Response to Tension Increase in the Tissue. Replot of data as in Figure 6 displayed as grouped data scatter plot of individual values. Individual measurements (repeats from different experiments, n=3 for each GeneX-siRNA) are depicted by blue circles with measurement range indicated by a connecting green line and the mean of three independent measurements by a black dash. Dark and light grey areas indicate the confidence level of 60 and 90%, respectively. Protein names are in black bold font for confidence levels above 90%, in black font for levels between 60 and 90 % and grey font for levels below 60%. Only samples whose % of breakage is below 5% are analyzed using this classification method (Figure 3).

